# Blockade of LAG-3 in PD-L1-deficient mice enhances clearance of blood stage malaria independent of humoral responses

**DOI:** 10.1101/2020.06.22.164566

**Authors:** Raquel Furtado, Laurent Chorro, Natalie Zimmerman, Erik Guillen, Emily Spaulding, Shu Shien Chin, Johanna P. Daily, Grégoire Lauvau

## Abstract

T cells expressing high levels of inhibitory receptors such as PD-1 and LAG-3 are a hallmark of chronic infections and cancer. Checkpoint blockade therapies targeting these receptors have been largely validated as promising strategies to restore exhausted T cell functions and clearance of chronic infections and tumors. The inability to develop long-term natural immunity in malaria-infected patients has been proposed to be at least partially accounted for by sustained expression of high levels of inhibitory receptors on T and B lymphocytes. While blockade or lack of PD-1/PD-L1 and/or LAG-3 was reported to promote better clearance of *Plasmodium* parasites in mice, how exactly these pathways contributes to protection is not known. Herein, using a mouse model of non-lethal *P. yoelii (Py)* infection, we reveal that the kinetics of blood parasitemia is indistinguishable between PD-1^-/-^, PD-L1^-/-^ and WT mice. Yet, monoclonal antibody (mAb) blockade of LAG-3 in PD-L1^-/-^ mice promoted accelerated control of blood parasite growth and clearance. We also report that i) the majority of LAG-3^+^ cells are T cells, ii) selective depletion of CD8^+^ T cells did not prevent anti-LAG-3-mediated protection, and iii) production of effector cytokines by CD4^+^ T cells is increased in anti-LAG-3-treated versus control mice. In addition, parasite-specific Ab serum titers and their ability to transfer protection from both groups of mice was comparable and depletion of CD4^+^ T cells prevented protection. Thus, taken together, these results are consistent with a model in which disruption of PD-L1 and LAG-3 on parasite-specific CD4^+^ T cells unleashes their ability to effectively clear blood parasites, independently from humoral responses.

**Author Summary:** Malaria, caused by *Plasmodium* parasites, is a global burden for which an efficacious vaccine is urgently needed. The development of long-term immunity against malaria is unclear, but we know that both T and B (that produce antibodies, Ab) lymphocytes, that are subsets of white blood cells, are required. Studies in mouse models of malaria have suggested that sets of inhibitory receptors, namely LAG-3 and PD-1, expressed on cytotoxic and helper T lymphocytes hamper the development of effective immunity against malaria. Therapeutic blockade of these receptors was reported to enhance blood parasite clearance through the development of more protective parasite-specific helper T lymphocytes and Abs. Herein, we reveal that, while mice genetically deficient for the PD-1 pathway fail to clear blood parasites better than WT counterparts, anti-LAG-3 treatment does. Importantly, we found comparable parasite-specific Ab responses between all mouse groups, and Ab transfers conferred similar protection to newly infected mice. We also show that LAG-3 is mostly expressed on T lymphocytes, and that cytotoxic T lymphocytes are not involved in anti-LAG-3 accelerated clearance of parasites. Our results suggest that LAG-3 blockade acts on helper T lymphocytes to unleash their effector responses and enhance the control of blood-stage malaria, independently from parasite-specific Abs.

## Introduction

The remarkable success of anti-cancer checkpoint blockade therapies have provided formal proof of concept that reinvigorating T cells with their effector potential is a feasible and realistic approach to achieve more effective immune-based therapies of human diseases (1). Understanding signals that can modulate and shape T cell functions has established the importance of costimulatory and coinhibitory pathways in T cell biology (2). While co-stimulation is essential to prime fully functional T cells, expression of inhibitory receptors such as program cell death 1 (PD-1) and Lymphocyte Activation Gene 3 (LAG-3) (3) expressed on activated T cells, enables regulation and optimal memory formation. Expression of PD-1 facilitates the down regulation of activated T cell functions through interactions with its major ligands PD-L1 and PD-L2. LAG-3 is a CD4 homologue that binds MHC class II on antigen-presenting cells with higher affinity than CD4 and can act as a negative regulator of T cell functions. Recently, the fibrinogen like protein 1 (FGL-1) that is naturally expressed by liver cells and overexpressed by some tumor cells, was characterized as a new ligand for LAG-3 which upon binding can potently prevent T cell activation (4).

In the context of chronic viral infections (HIV, Hepatitis C) and in tumors, sustained antigenic stimulation and an immuno-suppressive microenvironment promote the onset of T cells that maintain high levels of PD-1 and LAG-3 and exhibit an “exhausted” or dysfunctional phenotype, which prevent robust host protective effector T cell responses (1, 5). Blockade of the PD-1/PD-L1 pathway can -at least partially-rescue T cell functionality and allow for virus and tumor clearance (6). The fundamental importance of targeting T cell inhibitory pathways for therapeutic purposes has been highlighted in various cancers (1). More recently, several reports have also provided strong evidence that these inhibitory pathways may account for the lack of long-term sterilizing immunity in malaria infected patients (7). Expression of high levels of PD-1 and LAG-3 on CD4^+^ and CD8^+^ T cells, and expansion of such cells amongst peripheral blood mononuclear cells (PBMCs) of malaria-infected patients has been reported (8, 9). In addition, transcripts encoding the butyrophilin family member butyrophilin-like 2 (BTNL2), another negative regulator of T-cell activation, were increased during malaria infection compared to convalescent uninfected controls (10). Likewise, a subset of “atypical” memory B cells (CD21^-^ CD27^-^CD10^−^), originally described as “exhausted” B cells in HIV- and HCV-infected patients (11, 12), are expanded in malaria patients and may contribute to the lack of effective long-lived immunity against this parasite (9, 11). Both PD-1 and LAG-3 inhibitory molecules are also upregulated in various surrogate non-lethal mouse models of blood stage malaria (8, 12, 13), and, as in chronic viral infections and tumors, their selective blockade accelerates pathogen elimination. While enhanced parasite clearance is accounted for by improved CD4^+^ T and B cell/antibody responses (8), other mechanisms involving the restoration of CD8^+^ T cell cytolytic functions are also proposed (13).

Because of the potential clinical significance of these inhibitory pathways during malaria infection, we investigated the relative contribution of PD-1/PD-L1, LAG-3 or both pathways in the clearance of *Plasmodium* parasites from the blood of infected mice, and how they modulate host immune responses to improve infection outcome. Using the non-lethal and non-chronic *P. yoelii* mouse model of blood stage malaria infection, we report that mice genetically deficient for PD-L1 exhibit comparable kinetics of blood parasitemia to WT counterparts. We also found that LAG-3 blockade in PD-L1^-/-^ mice similarly accelerates parasite clearance as WT mice co-treated with anti-LAG-3/PD-L1 monoclonal Abs (mAbs). Yet, while therapeutic blockade of PD-L1/LAG-3 in WT mice promotes a greater magnitude of CD4^+^ T_FH_ and GC B cell responses, that of LAG-3 in PD-L1^-/-^ mice fails to enhance these responses. Rather, we reveal that blockade of LAG-3, which is mostly detected on activated CD4^+^ and CD8^+^ T cells, promotes parasite clearance independent of parasite-specific Abs and CD8^+^ T cell responses. Since CD4^+^ T cells are required for blood parasite elimination, these results are consistent with a model in which blockade of LAG-3 and PD-L1 act synergistically on CD4^+^ T cells to mediate direct parasite clearance.

## Results

### Detailed characterization of the robust B cell and associated follicular helper CD4^+^ T cell responses induced by non-lethal *P. yoelii* infection

To characterize the immune response of mice infected with the non-lethal strain of *Plasmodium, P. yoelii 17XNL (Py)*, we measured blood parasitemia and B and CD4^+^ T_FH_ cell responses over a 30 day period of infection (Fig 1 and S1 Fig). The proportion of infected red blood cells (iRBC) increased to a maximum of ∼70% of the RBC by day 15-17 before it cleared to undetectable levels by ∼day 22 (Fig 1). GFP-expressing parasites, most likely in circulating RBC, were detected in the red pulp area of spleen sections from infected mice (S1A Fig). Consistent with prior studies (8, 14), this observation suggests that the immune mechanisms underlying the control of parasite growth in the blood of infected mice occur during the first 15 days of infection. Since B cells play an essential role in protective immunity against malaria (8, 15), we carried out an in depth kinetic analysis of this cellular compartment in the spleen of *Py*-infected mice at 7.5, 12.5 and 15.5 days post infection (Fig 1B and S1B Fig). Using advanced flow-cytometry gating strategies to define subsets of splenic B cells (16), we subdivided the CD19^+^ B cells based on IgD and IgM cell-surface expression. Remarkably, by 7.5 days and peaking at 12.5 days, ∼40% of CD19^+^ B cells underwent isotype switching, becoming IgM^low^IgD^low^, while proportions of all other B cell subpopulations decreased (S1B Fig). IgM^low^IgD^hi^ B cells modestly diminished over time post infection, and represented ∼50% of CD19^+^ B cells that include type I follicular B cells (FOL I). We also noted reduction of IgM^hi^IgD^low^ B cells, which likely converted to marginal zone (MZ), and included B1 B cells. All B cell transitional stages (T1, T2, T3) proportions decreased during infection, yet marginal zone precursors (MZPs) and type II follicular B cells (FOL II) were largely maintained. The most noticeable changes occurred in the germinal centers (GC, GL7^+^CD95^+^) representing ∼15-23% of splenic B cells between day 7.5 and 15.5 post infection (Fig 1C). Plasmablast generation (CD138^+^PNA^+^) quickly increased following *Py*-infection to peak by day 7.5, and represented ∼7-10% of the splenic B cells. Production of effective GC and plasma B cell responses requires CD4^+^ T_FH_ cell responses (17), and, as expected, a substantial proportion of T_FH_ (ICOS^+^Bcl6^+^) formed by 7.5 days post infection (∼30% of CD4^+^ T cells) while only few follicular regulatory T cells (T_FR,_ Foxp3^+^Bcl6^+^) were generated, representing ∼7% of Foxp3^+^ T cells and ∼0.5-1% of all CD4^+^ T cells (Fig 1D). Altogether these results provide a comprehensive kinetic analysis of the robust B cell and associated CD4^+^ T_FH_ cell responses that are induced during blood stage malaria infection, in the non-lethal *Py* mouse model.

**Fig 1.**
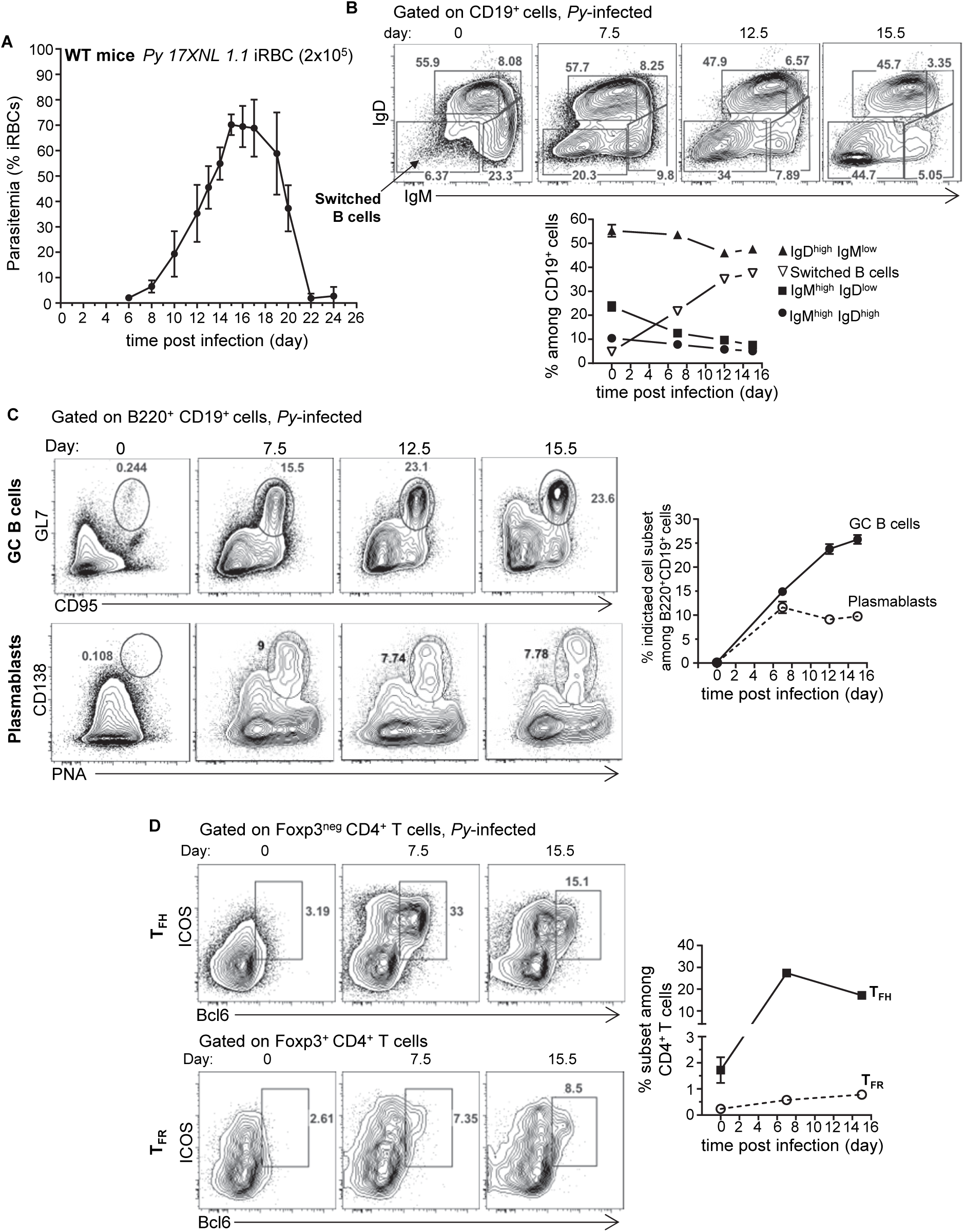
Kinetic analysis of B cell and follicular CD4^+^ T cell responses in *Plasmodium yoelii*-infected mice. WT B6 mice were inoculated with 2×10^5^ *Plasmodium yoelii (Py) 17XNL1.1* infected red blood cells (iRBC) i.v. **(A)** Blood parasitemia was measured starting 6 days post infection and every other day until day 24 using YOYO-1 staining of RBC and FACS. **(B-D)** Spleens from *Py*-infected mice were harvested at indicated days post infection and cells stained with mAb against lineage markers (CD19, B220, CD3, CD4, Foxp3, Bcl6) and activation/subset markers (IgD, IgM, CD93, CD23, CD21/35, GL7, CD138, CD95, PNA). In all experiments, representative FACS dot plots of 3-5 independent replicate experiments are presented (n= 5-18). Graphs show average results from experiments with SEM. P-values are indicated when applicable.

### Growth and clearance of blood parasites and adaptive immune responses in PD1^-/-^, PD-L1^-/-^ and heterozygous littermates is comparable to that of wild type mice

Overexpression of inhibitory molecules such as PD-1, LAG-3 or BTNL-2 on antigen-experienced T cells in malaria-infected patients were suggested as possible mechanisms accounting for the impaired development of effective and long-lasting adaptive immunity (7, 8, 10). To further assess the respective roles of these molecules in malaria, we inoculated *Py* iRBC in mice genetically deficient for the genes encoding for PD-1 or PD-L1 (PD1^-/-^, PD-L1^-/-^, namely KO), littermate counterparts possessing only one functional copy of either gene (PD1^+/-^, PD-L1^+/-^, namely heterozygotes) or both copies (PD1^+/+^, PD-L1^+/+^, namely WT) (Fig 2). The kinetics of *Py* growth and elimination was similar between all these groups (Fig 2A). Consistent with the parasitemia results, GC formation and plasmablast, CD4^+^ T_FH_ and T_FR_ cell differentiation at 7.5 and 13.5 days post infection, were also equivalent between all groups of mice (Fig 2B, 2C and S1C Fig).

**Fig 2.**
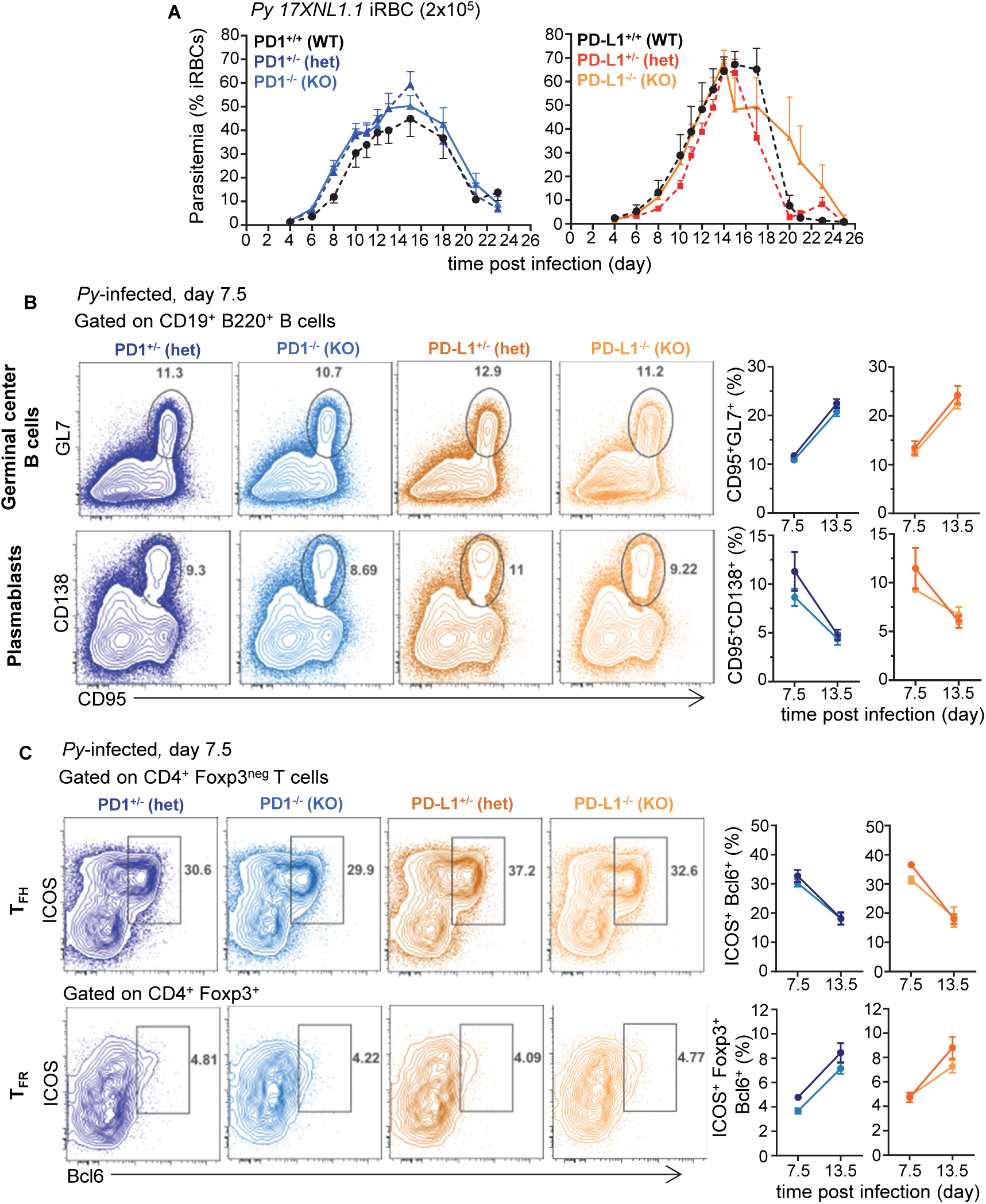
B and CD4^+^ T cell responses are comparable in *Plasmodium yoelii*-infected PD1^-/-^, PD-L1^-/-^, heterozygous counterparts and WT littermates. Littermates of indicated genotypes were inoculated with 2×10^5^ *Py 17XNL1.1* iRBC i.v**. (A)** Kinetics of *Py*-iRBC using YOYO-1 staining of RBC. **(B)** Spleens from *Py*-infected mice were harvested 7.5 and 13.5 days post infection and stained for CD19, B220, GL7, CD138, CD95 to monitor B cell responses and **(C)** spleen cells were stained for CD4, CD3, Foxp3, Bcl6, ICOS to monitor CD4^+^ T_FH_ and T_FR_ cellular responses by flow cytometry. In all experiments, representative FACS dot plots of 2 independent replicate experiments are presented (n=3-12). Graphs show average results from experiments with SEM. P-values are indicated when applicable.

We next assessed if T cells isolated from the various groups of *Py*-infected mice were functionally different compared to non-infected counterparts (Fig 3). Splenocytes from naïve or *Py*-infected WT mice were restimulated *ex vivo* (with PMA/ionomycin) and production of effector cytokines (IL-2, TNF, IFNγ) by CD4^+^ and CD8^+^ T cells was measured over a short time period (Fig 3A). The proportion of single and double cytokine-secreting T cells was significantly reduced compared to that of uninfected controls (by ∼75-90%). PD-1^-/-^ and PD-L1^-/-^ and heterozygous mice were also comparable and exhibited a loss of cytokine-secreting capacity as infection progressed (Fig 3C and S1D Fig). Thus, altogether, the lack of observed differences in i) blood parasite growth and elimination, ii) B and CD4^+^ T_FH_ cell responses and iii) production of effector cytokines across the distinct experimental groups, suggest a relatively modest contribution of the PD1/PD-L1 pathway in natural protection against blood stage malaria, a result that contrasts with prior published reports using PD-L1 Ab blockade in WT mice (8, 13).

**Fig 3.**
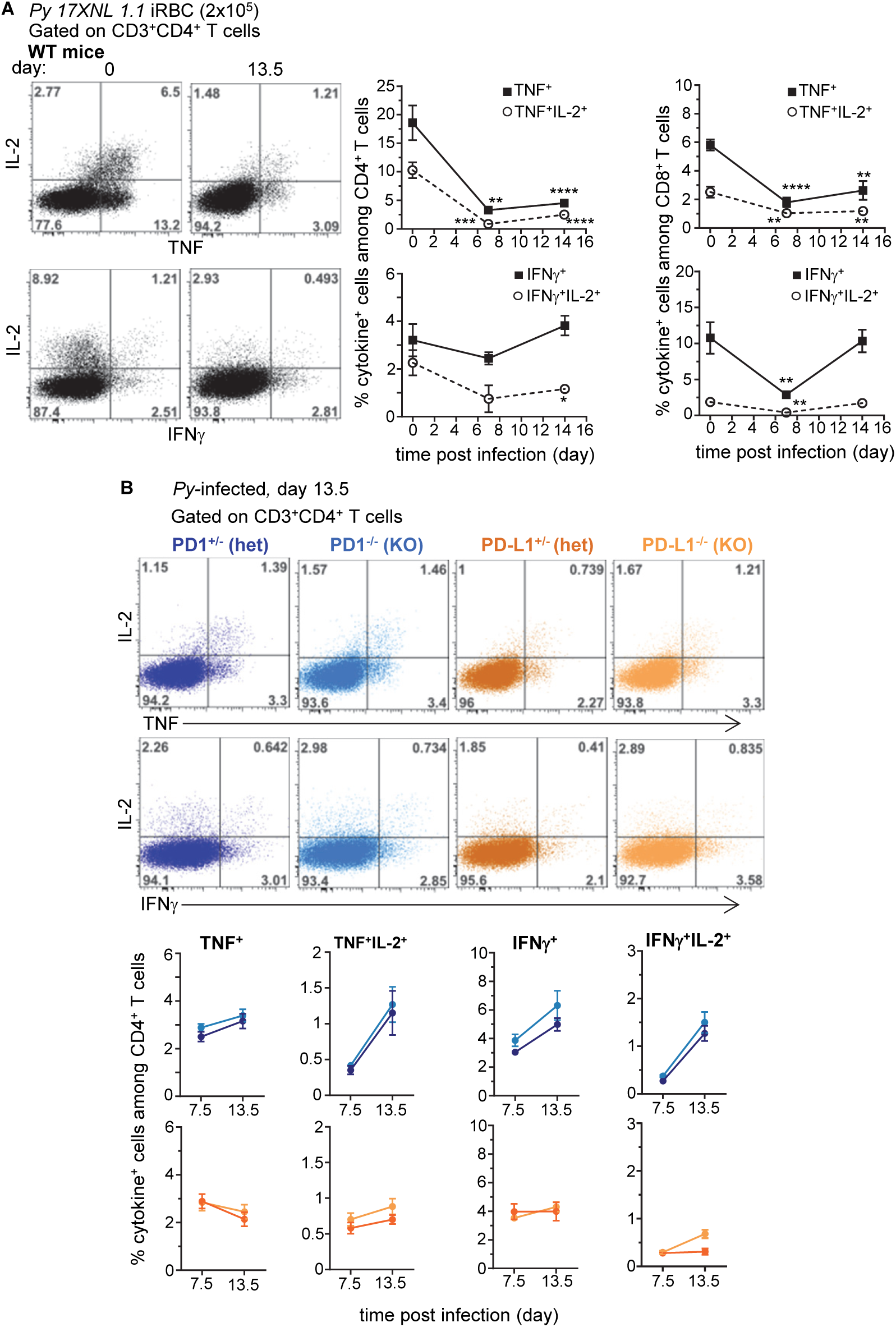
Dysfunctional T cell response in *Plasmodium yoelii*-infected WT, PD-1^-/-^ and PD-L1^-/-^ mice. Spleen cells from either uninfected, day 7.5 or 13.5 *Py*-infected (2×10^5^ iRBC, i.v.) **(A)** WT, **(B)** PD-1^-/-^ (KO), PD-1^+/-^ (hets), PD-L1^-/-^ (KO), PD-L1^+/-^ (hets) mice were stained for cell-surface CD3, CD4 and CD8 and intracellular cytokines IL-2, IFNγ and TNF. Frequencies of cytokine-producing cells among indicated T cell subset are shown. Graphs average 3-5 independent replicate experiments with SEM. P-values are indicated when applicable (*p<0.05, **p<0.01, ***p<0.001, ****P<0.0001).

### Blockade of LAG-3 promotes better control of blood parasite growth and clearance in PD-L1^-/-^ than in WT mice

Since therapeutic co-blockade of PD-L1 and LAG-3 inhibitory molecules in WT mice was reported to act synergistically to decrease blood parasite growth and accelerate its clearance (8), we hypothesized that LAG-3 blockade would promote more effective parasite elimination in PD-L1^-/-^ compared to WT mice. We injected WT and PD-L1^-/-^ mice with anti-LAG-3 and anti-LAG-3/PD-L1 or anti-LAG-3/PD-1 mAbs, respectively (Fig 4). As predicted (8), LAG-3 blockade in PD-L1^-/-^ mice promoted reduced parasitemia, suggesting faster control of blood parasite growth and elimination compared to WT counterparts, a result that we also confirmed in female mice S2A Fig). Treatment of PD-1^-/-^ mice with anti-LAG-3 mAb further confirmed that LAG-3/PD-L1 co-blockade in WT mice results in a synergistic enhancement of parasite elimination (S2B Fig). Moreover, therapeutic blockade of PD-1 in PD-L1^-/-^ mice, which neutralizes PD-1/PD-L2 interactions, resulted in delayed blood parasite clearance, consistent with a prior report (18) showing that blockade of the PD-1/PD-L2 axis prevents effective parasite control (S2C Fig). Interestingly, blood parasitemia was comparable in anti-LAG-3/PD-1-versus anti-LAG-3-treated PD-L1^-/-^ mice, suggesting that inhibition of LAG-3 is dominant over that of PD-1/PD-L2. Collectively, these findings i) confirm the importance of PD-1-dependent and LAG-3 inhibitory pathways in resistance to malaria infection and ii) reveal that blockade of a specific inhibitory pathway, here LAG-3, can lead to distinct outcomes depending on genetic or functional deficiencies in other related pathways, here PD-L1 or PD-L2.

**Fig 4.**
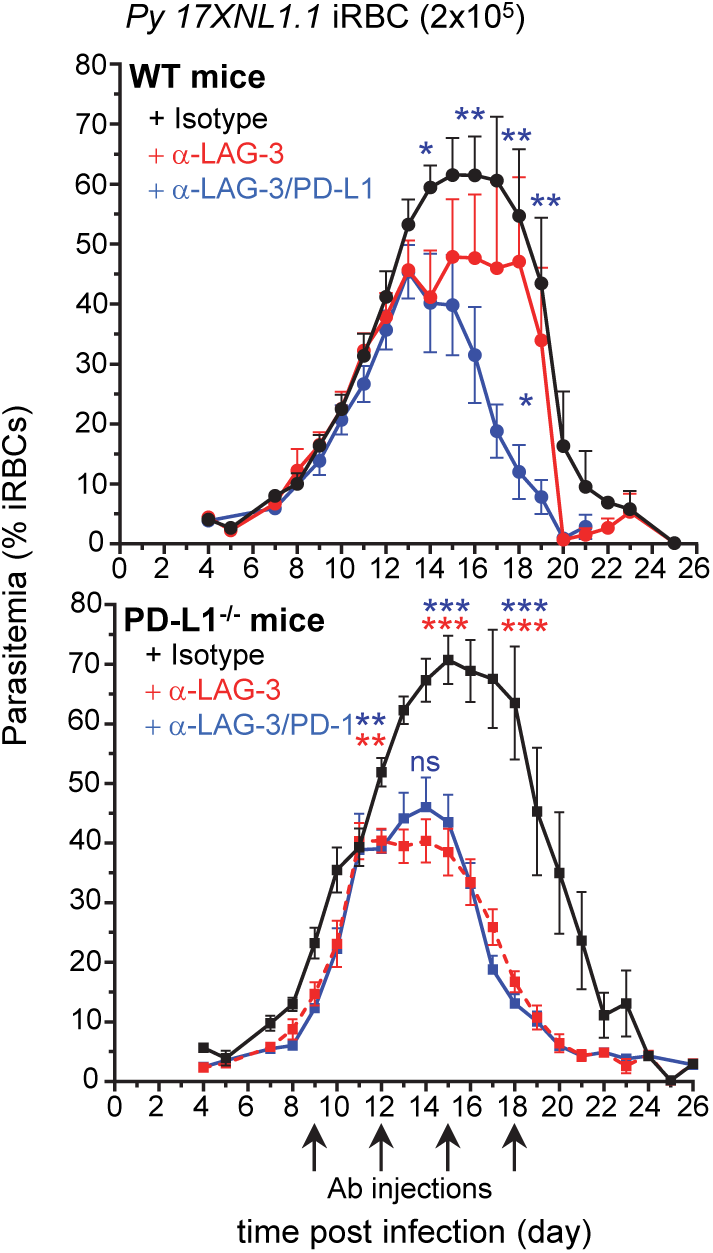
LAG-3 blockade enhances blood parasite clearance. WT or PD-L1^-/-^ mice were inoculated with 2×10^5^ *Py 17XNL1.1* iRBC i.v., and 9, 12, 15 and 18 days post infection, indicated mice received 200 μg of either anti-LAG-3, anti-LAG-3/PD-L1, anti-LAG-3/PD-1 or polyclonal rat IgG Ab (isotype) i.v.. Results show the kinetics of blood parasitemia over time determined by YOYO-1 staining of RBC collected at indicated time points. Graphs average the pool of 2-3 independent experiments (n=10-13) shown with SEM. P-values are indicated when applicable (*p<0.05, **p<0.01, ***p<0.001) for each treatment compared to isotype (asterisk color indicate respective treatment group) or to anti-LAG3 treatment per time point (middle blue asterisk or “ns” on graphs for anti-LAG3/PD-L1 compared to anti-LAG3).

### Antigen-experienced T_bet_^+^ CD8^+^ and T_bet_^+^, T _FH_ and T _FR_ CD4^+^ T cells are the major LAG-3-expressing cell subsets

To gain further insight into how LAG-3 blockade contributes to reduced blood parasitemia, we next investigated which cells express the LAG-3 receptor during *Py* infection (Fig 5). LAG-3-expressing spleen cells represented ∼8.5% of live cells 7.5 days post-*Py* infection and largely consisted of T cells (>80%) that make up ∼26% of splenic cells, distributed among CD4^+^ and CD8^+^ T cells (Fig 5A and 5B). Using a 26 color flow cytometry panel which includes cell surface lineage and activation markers, intracellular functional markers and lineage specifying transcription factors, we found that activated and proliferating (CD62L^lo^CD44^hi^Ki67^+^) CD8^+^ T cells distributed into T_bet_^+^, Eomes^+^ and T_bet_^+^ /Eomes^+^ effector cells (Fig 5C and S3A Fig). T_bet_^+^ cells expressed high levels of the cytolytic marker granzyme B and cell-surface marker CX3CR1 which are features of terminally differentiated effector cells (19). Interestingly, activated and proliferating CD4^+^ T cells mostly included T_bet_^+^ and T_FH_ cells, both of which expressed high levels of the cell-surface marker ICOS (Fig 5D). In contrast, ICOS^neg^ CD4^+^ T cells were not activated (CD62L^hi^CD44^lo^Ki67^neg^) (S3B Fig). Among LAG-3^+^ CD4^+^ T cells, a majority were activated ICOS^+^ conventional T cells (∼48%), T_FH_ cells (∼33%) and T_FR_ cells (∼32%) (Fig. 5B). Some Foxp3^+^ T_reg_ cells also expressed LAG-3 but only represented a small proportion of all CD4^+^ T cells (<6%). Interestingly, using CD49d and CD11a cell-surface markers as a proxy for activated parasite-specific T cells, we found a substantial proportion of LAG-3^+^ T cells expressed both markers (∼84% and ∼38% respectively, for CD8^+^ and CD4^+^ T cells) and >90% also upregulated the adhesin CD11a (Fig 5E). A large proportion of CD49d^+^/CD11a^+^ CD8^+^ and CD4^+^ T cells were activated and proliferated (CD62L^lo^CD44^hi^Ki67^+^), and expressed T_bet_, Eomes and were T_FH_ (for CD4^+^ T cells) (Fig 5F). Thus, these data support a model in which the largest proportion of LAG-3^+^ cells are T cells that are likely parasite-specific, and include effector T_bet_^+^ CD8^+^ T cells and T_bet_^+^, T_FH_ and T_FR_ CD4^+^ T cells.

**Fig 5.**
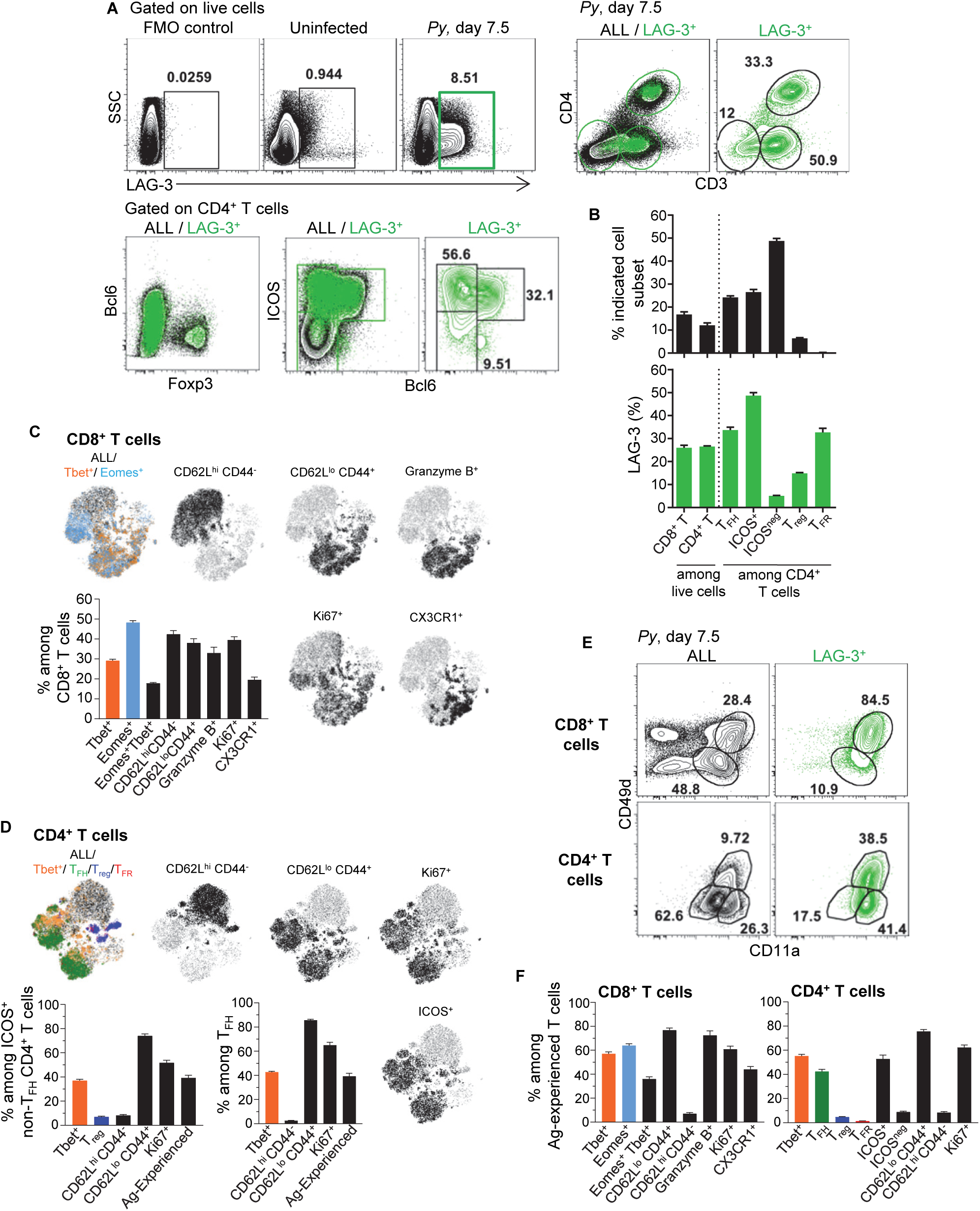
LAG-3 is expressed on activated CD4^+^ and CD8^+^ T cells during *Py* infection. Spleen cells from day 7.5 *Py*-infected mice were stained for lineage markers (CD8, CD4, CD3, Foxp3, Bcl6) and activation/inhibitory markers (LAG-3, CD11a, CD49d), or with a 26 color panel in S3A Fig (C, D, F). **(A)** FACS dot plots of the phenotype and **(B)** proportion of LAG-3-expressing cells among the indicated T cell subsets. **(C, D)** t-SNE overlay representation of indicated CD8^+^ (C) and CD4^+^ (D) T cell subsets from a pool of 5 infected WT mice with bar graphs summarizing the proportions of indicated subsets with SEM. **(E)** Proportion of Ag-experienced (CD11a^+^CD49d^+^) CD8^+^ and CD4^+^ T cells on all or LAG-3^+^ cells. **(F)** Proportions of indicated subsets among Ag-experienced CD8^+^ and CD4^+^ T cells from the pool of 5 infected WT mice with SEM. Overall, representative FACS dot plots of 2 independent replicate experiments are presented for 1 of 3-5 mice. Graphs show average results from experiments with SEM. P-values are indicated when applicable.

### Anti-LAG-3-mediated parasite elimination is independent from CD8^+^ T cells

Since the major LAG-3^+^ cell types are CD8^+^ and CD4^+^ T lymphocytes, we next tested if selective depletion of either T cell subset would abrogate the LAG-3-mediated protective effect (Fig 6). PD-L1^-/-^ mice were either injected with anti-CD8β or anti-CD4 depleting mAbs or PBS prior to infection with *Py*, and 9, 12 and 15 days post-infection, mice received anti-LAG-3 or control isotype Abs. We monitored weight and parasitemia every other day until the undepleted isotype Ab-treated control groups were negative for blood parasitemia. LAG-3 blockade in CD8^+^ T cell-depleted PD-L1^-/-^ mice enhanced the control and clearance of blood parasitemia similarly to that of undepleted control groups, demonstrating that anti-LAG-3 blockade does not act on CD8^+^ T cells. In contrast, CD4^+^ T cell-depleted mice failed to control parasite growth and developed chronic parasitemia, and lost >10% of initial body weight 20 days post infection whether treated with anti-LAG-3 or control isotype Abs, consistent with their requirement for *Py* parasite clearance (8). Taken together with the finding that LAG-3 is mostly expressed on T cells (Fig 5), these results suggest that the anti-LAG-3-mediated therapeutic effect on blood parasite elimination is likely to occur by blocking LAG-3 on parasite-specific T_bet_^+^ and/or T_FH_ CD4^+^ T cells, but not CD8^+^ T cells.

**Fig 6.**
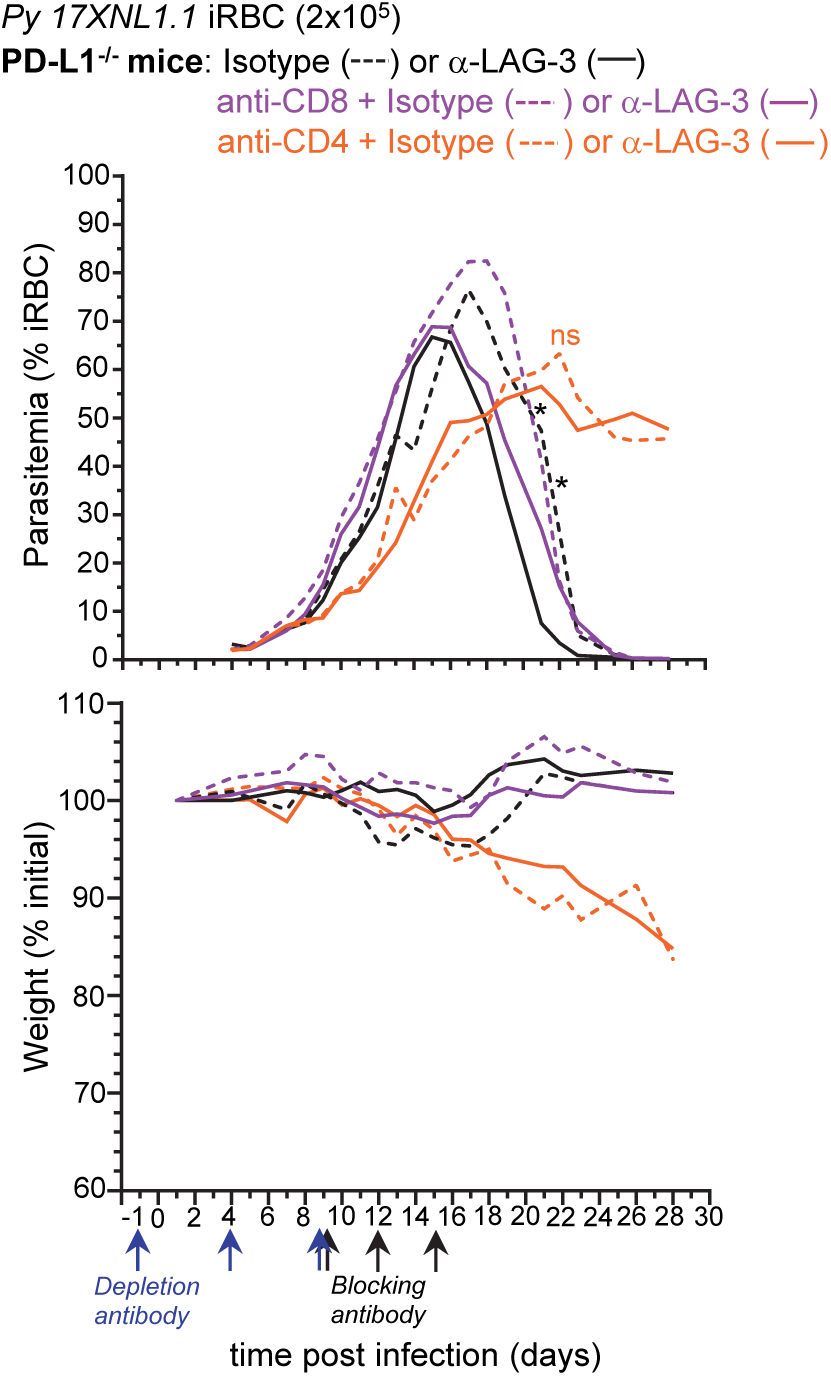
LAG-3-mediated enhancement of anti-parasite immunity is not dependent on CD8^+^ T cells. PD-L1^-/-^ mice were injected i.p. with PBS or 150µg of either anti-CD4 or anti-CD8β depleting mAbs as depicted and inoculated with 2×10^5^ *Py 17XNL1.1* iRBC i.v. Mice also received either 300 µg anti-LAG-3 or isotype Ab i.v. on day 9, 12 and 15 post *Py* infection. Results show blood parasitemia measured by YOYO-1 staining of RBC, and change in initial weight (lower panel) over time starting on day 4 post *Py* infection. Graphs are representative of two independent experiments (n=3-4) shown with SEM. P-values are indicated when applicable (*p<0.05) for each treatment compared to isotype (asterisk color indicate respective group)

### LAG-3 neutralization in PD-L1^-/-^ mice fails to promote the production of higher titers and more protective parasite-specific antibodies

Since the reported mechanism by which LAG-3/PD-L1 blockade in WT mice enhances parasite clearance involves greater GC B and CD4^+^ T_FH_ cell responses and higher titers of parasite-specific Abs (8), we next hypothesized that LAG-3 neutralization in PD-L1^-/-^ mice may work through a similar mechanism. We monitored both CD4^+^ T_FH_ and B cell GC and plasmablast responses in PD-L1^-/-^ and WT mice treated with anti-LAG-3 or anti-LAG-3/PD-L1, respectively (Fig 7). While as expected, blockade of LAG-3 and PD-L1 in WT mice promoted significantly enhanced T_FH_ and GC B cell responses, and compared to uninfected mice, we did not find any differences in these cell populations in anti-LAG-3-treated PD-L1^-/-^ mice. The proportion of switched B cells was also comparable in all experimental groups (not shown). Of note, spleen weight and total or hematopoietic-derived (CD45^+^) cell counts per spleen were comparable in WT and PD-L1^-/-^ mice undergoing the various treatments, likely ruling out that the lack of differences in cell subset proportions in the various experimental groups is accounted for by differences in absolute cell numbers (S2D Fig).

**Fig 7.**
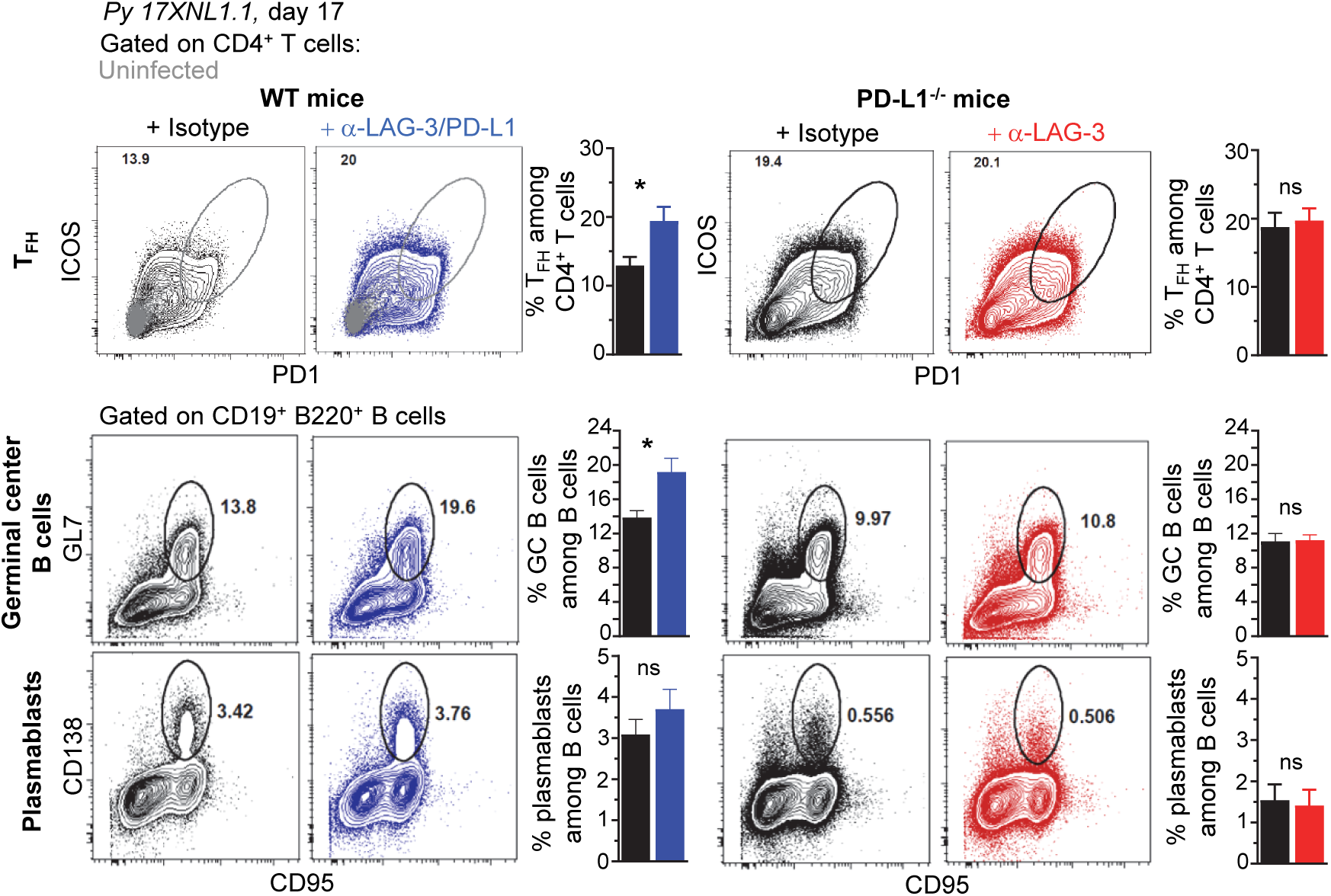
LAG-3/PD-L1 therapeutic blockade enhances T_FH_ and GC responses while LAG-3 blockade in PD-L1^-/-^ mice does not. WT B6 or PD-L1^-/-^ mice were inoculated with 2×10^5^ *Py 17XNL1.1* iRBC i.v. and 9, 12, 15 and 18 days post infection, indicated mice either received 200 μg of either anti-LAG-3/PD-L1, anti-LAG-3 or isotype Ab i.v. Spleens were harvested 17 days post infection and stained for CD4, CD3, ICOS, PD-1 to monitor CD4^+^ T_FH_ cell response (ICOS^+^PD-1^hi^ corresponds to ICOS^+^Bcl6^hi^ cells) (top panel) and for CD19, B220, GL7, CD138, CD95 to monitor B cell responses (lower panels). Representative FACS dot plots of 1-2 independent replicate experiments are presented (n=4-10), with an overlay of representative uninfected mouse in the WT top panel. Graphs average each experiment shown with SEM. P-values are indicated when applicable (*p<0.05).

We next conducted experiments to assess if, despite the lack of obvious differences in the B and CD4^+^ T_FH_ cell responses in PD-L1^-/-^ mice, parasite-specific Abs produced in anti-LAG-3-treated group may still confer better protection and exhibit higher serum titers compared to isotype-treated counterparts (Fig 8). We transferred sera (150 μl/mouse, one time) isolated from anti-LAG-3 or isotype Ab-treated PD-L1^-/-^ mice 18, 32 or 72 days post *Py* infection, into WT B6 recipient mice that were subsequently infected with *Py*. Data reveal a delayed onset and peak blood parasitemia, and lower weight loss in mice treated with day 72 sera, but no beneficial trends were observed in day 18 or 32 sera transfers. This result was extended in sera isolated from day 32 *Py*-infected WT mice treated with anti-LAG-3/PD-L1 or isotype control Abs, suggesting that, at least under these experimental conditions, transfer of parasite-specific Abs from *Py*-infected mice failed to confer significantly distinct levels of protection to recipient mice (S4A Fig). Yet, immune sera from day 72-infected WT mice did confer increased protection and prevented weight loss compared to pre-immune sera, validating our serum transfer experiments and read-outs. Along these lines, only sera from day 72 anti-LAG-3, but not isotype-treated infected PD-L1^-/-^ mice, delayed parasitemia in recipient mice that were subsequently infected, suggesting that PD-L1^-/-^ mice are unable to develop long-term parasite-specific protective Abs (Fig 8). Comparative day 32 sera transfers between isotype Ab-treated WT and PD-L1^-/-^ mice showed no differences in parasitemia in recipient mice, yet slightly better *Py* clearance and lower weight loss for day 72 sera from WT mice was observed compared to that of PD-L1^-/-^ mice (S4 Fig). This further supported the idea that PD-L1^-/-^ mice do not develop effective parasite-specific protective Abs.

**Fig 8.**
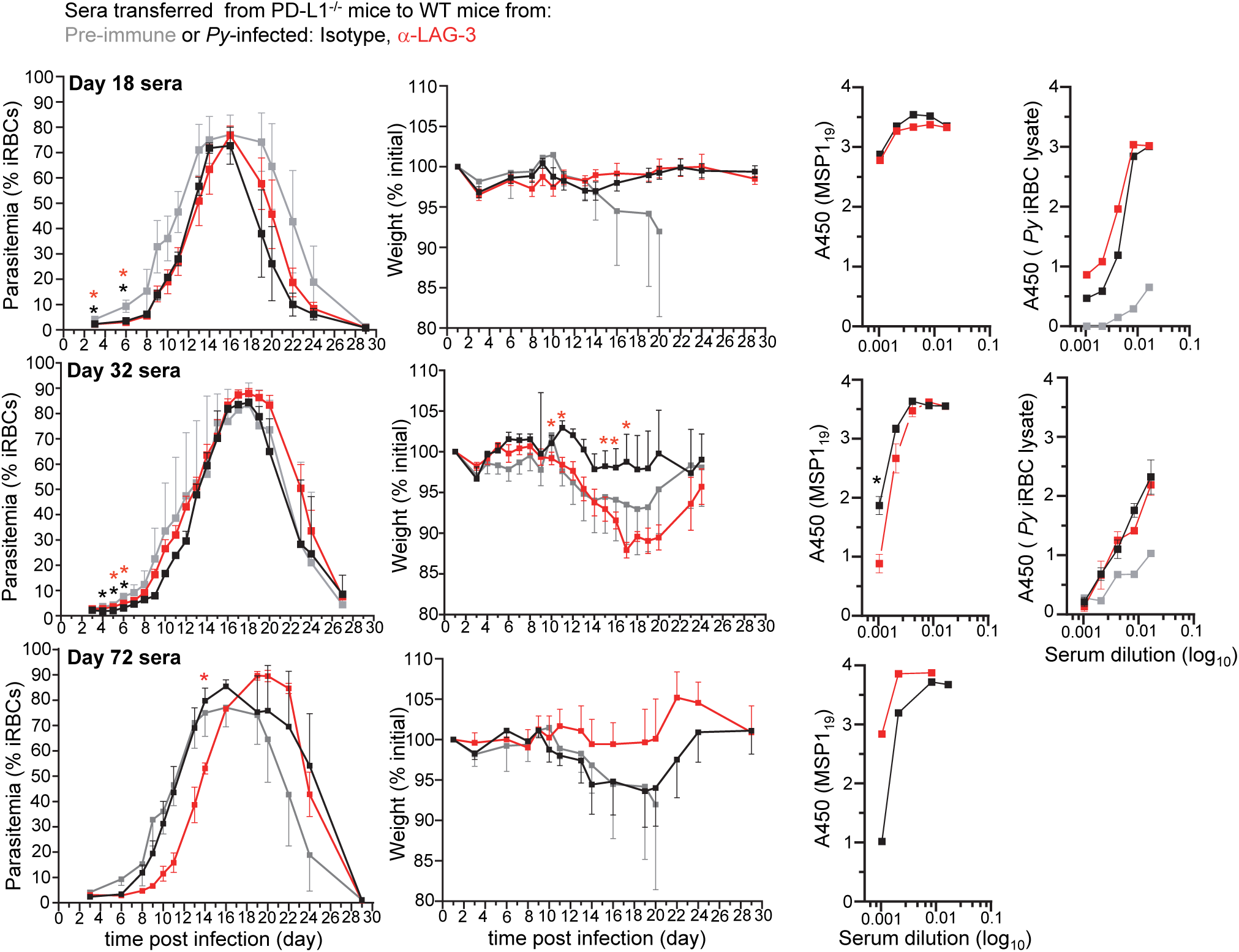
Transfer of parasite-specific serum Abs in anti-LAG-3-treated PD-L1^-/-^ mice do not confer better protection than control groups and all *Py*-specific Ab titers are comparable. 150 μL of sera harvested from mice 18, 32 or 72 days post *Py*-infection was transferred to naïve WT mice (3-5 mice/group) that were subsequently infected with 2×10^5^ *Py 17XNL1.1* iRBC i.v. Blood parasitemia and weight was monitored over 28 days. Titers of *Py* MSP1_19_-specific or *Py* iRBC-specific IgG antibodies in indicated sera was measured by ELISA and reported as background subtracted A_450_ values. P-values are indicated when possible compared to pre-immune treatment or to isotype treatment. Asterisk color indicate respective treatment group (*p<0.05).

To provide additional evidence to these findings, we also measured parasite-specific Ab titers using i) recombinant merozoite surface antigen MSP1_19_, the major blood stage parasite Ag and ii) a lysate of *Py* iRBCs (Fig 8 and S4 Fig). Consistent with the serum transfer experiments, titers of parasite-specific Abs specific for both MSP1_19_ and *Py* iRBC extracts in the sera of anti-LAG-3 versus isotype Ab-treated PD-L1^-/-^ mice did not reveal significant differences. Slightly higher Ab titers were nevertheless measured in day 72 sera from anti-LAG-3-treated PD-L1^-/-^ mice, in line with the observed delayed parasitemia results (Fig 8). Likewise, slightly higher parasite-specific Ab titers were quantified in day 72 sera from anti-LAG-3/PD-L1-treated WT mice (S4A Fig).

In conclusion, these results suggest that the mechanism of LAG-3 blockade in PD-L1^-/-^ mice and LAG-3/PD-L1 co-blockade in WT mice that allows rapid control of blood parasitemia is unlikely to involve the production of higher titers of parasite-specific Abs. Also consistent with prior studies, our data show that peak production of such protective Abs most likely occurs within 2-3 months after infection, but not at early stages (20).

### CD4^+^ T cells from anti-LAG-3-treated PD-L1^-/-^ mice produce higher levels of effector cytokines than untreated counterparts

Our results so far have led us to hypothesize that blockade/disruption of the LAG-3 and PD-1/PD-L1 pathways enhances rapid parasite clearance via CD4^+^ T cells, independent from humoral responses. Since therapeutic blockade of these pathways was also shown to restore malaria-induced T cell dysfunction (8, 9), we next tested if T cells isolated from LAG-3-treated PD-L1^-/-^ mice were more responsive to broad PMA/ionomycin stimulation than control groups at day 17, when rapid parasite elimination is observed (Fig 9). We report significant differences by a factor of 2-3 fold, in the relative proportion and numbers of single (IL-2^+^) and double (IL-2/TNF^+^, IL-2/IFNγ^+^, IFNγ/TNF^+^) cytokine-secreting CD4^+^, but not CD8^+^, T cells in the anti-LAG-3-treated group compared to the control group, consistent with our hypothesis (Fig 9A and 9B). While a higher proportion of anti-LAG-3-treated CD4^+^ T cells secreted effector and proliferative cytokines, the proportion of the various phenotypic and functional subsets of CD4^+^ and CD8^+^ T cells (T_FH_, T_reg_, T_FR_, Ag-experienced) remained similar between all experimental groups and conditions (Fig 9C and 7A). High dimensional flow-cytometry analysis of the CD4^+^ T cells at later stages (day 36) using a 25 color panel (S3A Fig), also failed to further reveal any CD4^+^ T cell or memory (CD62L^+^ CD44^lo^) CD4^+^ T cell subset differences between anti-LAG-3 and the control isotype Ab-treated PD-L1^-/-^mice, confirming our findings (Fig 9D and S3C Fig). In summary, LAG-3 blockade contributes to rescuing malaria-induced CD4^+^ T cell dysfunction in PD-L1^-/-^ mice, but does not appear to modulate their ability to differentiate into various functional subsets of T cells.

**Fig 9.**
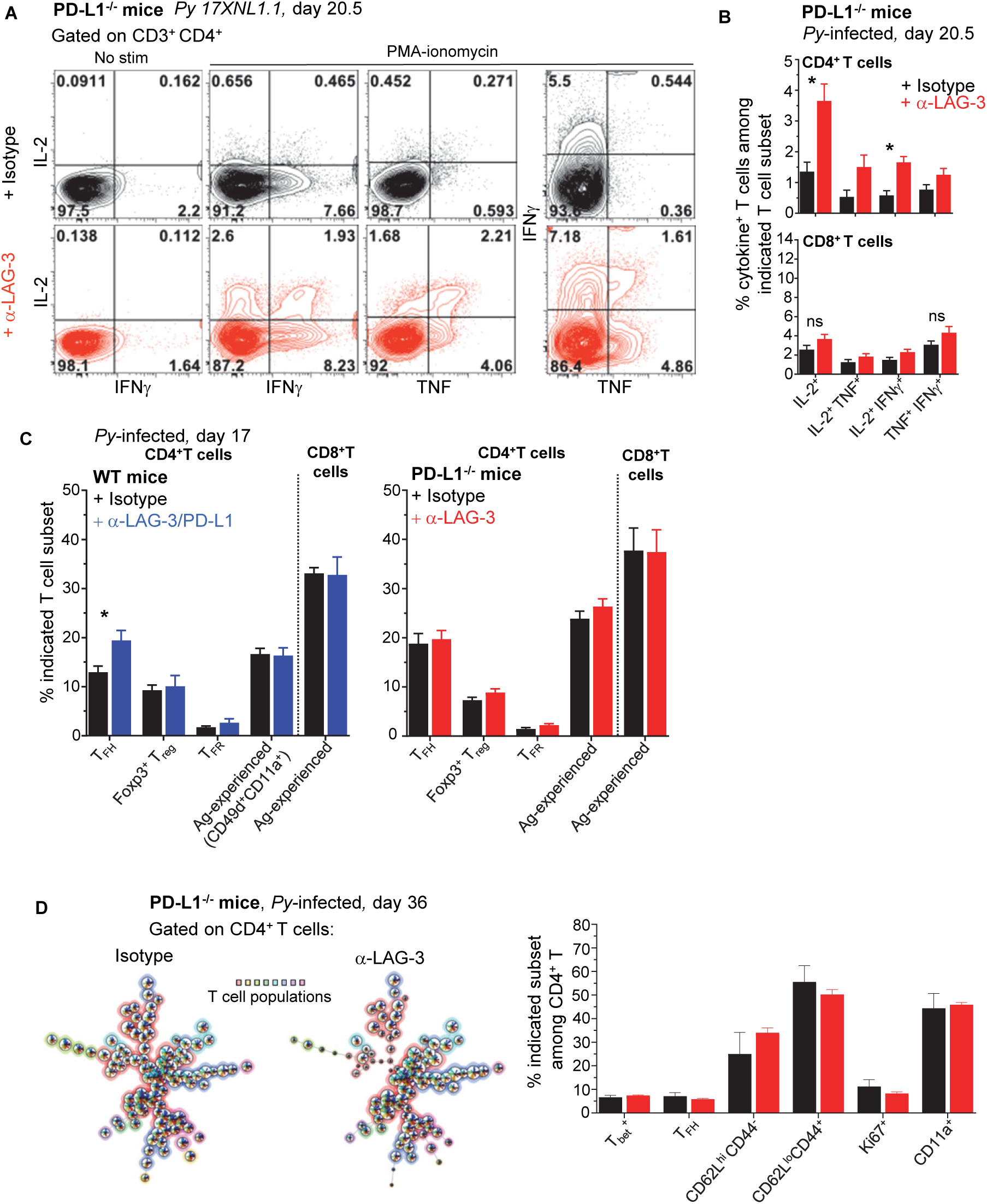
LAG-3 blockade in PD-L1^-/-^ mice partially restores CD4^+^ T cell dysfunction. Spleen cells from day 20.5 *Py*-infected PD-L1^-/-^ or WT mice, treated with either anti-LAG-3/PD-L1, anti-LAG-3 or isotype Ab i.v. (as indicated, on days 9, 12, 15) were stained for cell-surface CD3, CD4 and CD8, Foxp3, Bcl6, ICOS and intracellular cytokines IL-2, IFNγ and TNF after incubation with PMA/ionomycin for 4 hours (T cell mix). **(A)** Representative FACS dot plots of 2 independent experiments are shown with the frequencies of cytokine-producing cells among CD4^+^ T cells (n=3-5). **(B)** Graphs of the average frequencies of cytokine-producing cells among CD4^+^ T cells (shown in (A)) and CD8^+^ T cells with SEM. P-values are indicated when applicable. **(C)** Distribution of known splenic T cell subsets as indicated on day 17 post infection, T_FH_ data are from the same experiment shown in Fig 7. Graphs average 1-2 independent experiments (n=4-5) shown with SEM. **(D)** Spleen cells from day 36 PD-L1^-/-^ *Py*-infected mice treated with either anti-LAG-3 or isotype Ab i.v. (on days 9, 12, 15) were stained for lineage markers (CD8, CD4, CD3, Bcl6, T_bet_) and functional markers (CD62L, CD44, CD11a) with a 24 color panel depicted in S3A Fig. FlowSOM analysis (left) on pooled CD4^+^ T cells (n=5) and summary bar graphs from each treatment group are shown (right). Distribution of known splenic T cell subsets as indicated on day 36 post infection.

## Discussion

In this work, we investigated the mechanisms by which blockade of the inhibitory receptors LAG-3 and PD-L1 promote more effective control and clearance of non-lethal blood stage murine *P. yoelii* malaria. We reveal that LAG-3 neutralization in mice that are genetically deficient for PD-1/PD-L1 do not promote stronger CD4^+^ T_FH_ and GC B cell responses as was reported in anti-PD-L1/LAG-3 treated WT mice (8). Consistent with this result, convalescent sera (day 32) transfers from anti-LAG-3 treated PD-L1^-/-^ mice and anti-PD-L1/LAG-3 treated WT compared to control isotype Ab treated mice failed to confer distinct levels of protection against *Py* infection in naïve recipient mice. Additionally, serum titers of parasite-specific Abs were comparable in all immunized groups of mice. We further found that LAG-3, at the time of Ab blockade, is mostly expressed by T cells, with a large majority consisting of activated proliferating Ag-experienced (CD49d^+^CD11a^+^) T cells that include T_bet_^+^ CD8^+^ and CD4^+^ T cells, and T_FH_ and T_FR_ CD4^+^ T cells. We show that, while LAG-3 is expressed by both subsets of T cells, the LAG-3-mediated protective effect does not require CD8^+^ T cells, thus is most likely to act on CD4^+^ T cells. We also found that LAG-3 blockade enhances CD4^+^ T cell effector functions in infected mice. Thus, we propose a model in which inhibition of LAG-3 on parasite-specific CD4^+^ T cells unleashes their ability to produce effector cytokines, altogether contributing to achieve superior levels of protection in malaria-infected hosts, independent from parasite-specific humoral responses.

Our finding that mice with genetic deficiency in the PD-1/PD-L1 pathway are as susceptible to *Py* malaria infection as their WT counterparts highlights some unexpected discrepancies with prior work showing that therapeutic blockade of PD-L1 in WT mice promotes enhanced control and clearance of blood parasitemia (8, 13). The fact that i) the kinetics of infection was comparable in *Py*-infected PD-1^-/-^, PD-L1^-/-^, heterozygous and WT littermate mice, and ii) we did not find any differences in either T, B cell or Ab responses despite extensive analyses, suggests that genetic deficiency in PD-1 and PD-L1 may favor the utilization of distinct mechanisms of resistance against this infection through otherwise redundant pathways. The indistinguishable kinetics of parasitemia in PD-1^-/-^ and PD-L1^-/-^ mice together with the LAG-3 blockade experiment support such an interpretation, since LAG-3 neutralization in PD-L1^-/-^ mice promotes remarkable growth control and clearance of blood parasites, which is only achieved in WT mice upon co-blockade with PD-L1. A similar observation has been reported in a model of ovarian cancer in which therapeutic co-blockade of PD-1/LAG-3/CTLA-4 in WT mice leads to two times fewer tumor free mice compared to anti-LAG-3/CTLA-4 treatment in PD-1^-/-^ mice (21). These results further illustrate a more generally accepted dogma that mice genetically deficient in important molecules often use compensatory defense mechanisms that rely on other redundant pathways. Thus, and more generally, it is conceivable that any genetic deficiency for specific inhibitory pathways is likely to alter the outcomes of specific therapies targeting these important pathways in immune defense and homeostasis.

While the genetic deficiency in PD-L1 versus therapeutic mAb blockade of PD-L1 in WT mice clearly results in different outcomes for *Py* parasitemia and immune responses, our experiments using anti-LAG-3 mAb blockade raises an important point related to the underlying mechanisms of increased resistance to *Py* infection. The Butler study reported significantly stronger T_FH_, GC and plasmablast responses, as well as higher titers of MSP1_19_ parasite-specific Abs in anti-PD-L1/LAG-3-treated WT mice compared to control groups. Whereas we did confirm enhanced T_FH_ cell and GC responses in anti-PD-L1/LAG-3 treated WT mice, this neither translated into higher parasite-specific Ab titers (both against MSP1_19_ and whole *Py* iRBC lysate), nor into a better ability to confer protection to sera-transferred recipient mice infected with *Py*. We extended these findings to anti-LAG-3 versus control isotype Ab treated PD-L1^-/-^ mice, suggesting that parasite-specific Ab responses are unlikely to account for the enhanced anti-parasite responses that follow LAG-3 mAb blockade. While our serum transfer experiments were conducted according to the Butler study, i.e., single transfer of 150 μL of pooled sera from convalescent mice (day 32) prior to *Py* infection of recipient mice, most previous studies showing significant protection against malaria upon serum transfer, used 200 μL and at least 3 serial transfers at the beginning of infection (22, 23). We also transferred sera isolated from mice at different time points post *Py* infection (day 18, 32 and 72) following the rationale that, if Abs are indeed mediated the more effective anti-parasite response, this effect should be even clearer at earlier times when the differences between anti-LAG-3 and control isotype Ab treated mice are the greatest (day 18). Yet our results did not support this interpretation. While we observe that sera from day 72 convalescent mice -a time point at which peak parasite specific Abs are produced (20)-conferred better protection to recipient mice compared to pre-immune sera, we did not find i) any significant differences in the ability to confer parasite-specific protection or ii) higher parasite-specific Ab titers in anti-LAG-3 or anti-PD-L1/LAG-3 treated PD-L1^-/-^ or WT mice, respectively. We also note that our anti-LAG-3 treatment regimen was only 4 consecutive injections, compared to 7 in the Butler study. While it is possible that this accounts for some of the differences in the outcomes, the day 32 sera transfers conducted in this prior work seem unlikely to fully reflect the protective mechanisms at work during anti-LAG-3 treatment between day 9 and 18 post *Py* infection. Taken together, we favor the interpretation that LAG-3 neutralization leads to improved control of parasitemia by rescuing more functional effector CD4^+^ T cell responses, independent of parasite-specific Ab responses. A high proportion of activated proliferating (CD62L^lo^CD44^hi^Ki67^+^) Ag-specific (CD49d/CD11a^+^) effector (T_bet_^hi^) CD4^+^ T cells are observed among LAG-3^+^ CD4^+^ T cells. We found that a greater frequency of these cells secrete effector cytokines (IFNγ, TNFα, IL-2) in anti-LAG-3 versus control mice. These effector cytokine-secreting CD4^+^ T cells may also promote the influx and activation of higher numbers of innate inflammatory cells, e.g., monocytes, macrophages and neutrophils, to the spleen where blood filtration takes place (24, 25).

We provide strong evidence that while both CD8^+^ and CD4^+^ T cells express LAG-3 at the time of anti-LAG-3 treatment (day 7.5), and represent the major LAG-3-expressing cell compartment during *Py* infection (>80%), LAG-3 blockade is most likely acting on CD4^+^ T cells to unleash specific key effector mechanisms that help induce a more effective anti-parasite immune response. Yet, despite an extensive analysis of these antigen-experienced, parasite-specific CD4^+^ T cells, across various time points post *Py* infection (day 7.5, 17 and 36) using high dimensional 26 color flow cytometry panels, t-SNE and FlowSOM analysis of the CD4^+^ T cells in anti-LAG-3 or control isotype-treated PD-L1^-/-^ mice, we did not find any differences in CD4^+^ T cell subsets. The proportion of activated and proliferating CD4^+^ T cells, and of T_bet_^+^ (Th1), T_FH_ and T_FR_ remained equivalent in all groups, consistent with the lack of detectable differences in the humoral response. Since cytokine production by the CD4^+^ T cells upon *in vitro* restimulation was the only functional read out that we found significantly different, we propose that LAG-3 blockade is likely to act as a rapid and likely temporary boost of activated CD4^+^ T cells in this infection. Perhaps similar to this experimental situation, anti-PD-1 treatment in chronic virus-infected mice re-invigorate exhausted CD8^+^ T cells, yet still fails to achieve long-term re-programming of these cells (26).

Even though our data support a functional role for CD4^+^ T cells in effective anti-parasite immunity, the fibrinogen-like protein 1 (FGL-1) appears as a major ligand of LAG-3 that works independently of MHC-II, and inhibits T cell-mediated anti-tumor immunity (4). While it is unknown whether levels of FGL-1 increase during malaria infection, an appealing hypothesis may be that FGL-1 contributes to inhibiting LAG-3 expressing subsets of T cells during infection, including CD8^+^ T cells. Along this hypothesis, we found a slightly delayed blood parasitemia clearance in anti-LAG-3-treated, CD8^+^ T cell-depleted PD-L1^-/-^ mice, and a mild increase in cytokines secreted after *ex vivo* stimulation of CD8^+^ T cells isolated from anti-LAG-3-treated *Py*-infected mice. In the *P. chabaudi* murine model of infection that induces low chronic blood-parasite rebounds over 1-3 months post infection (14), CD8^+^ T cells were reported to be essential in mediating effective parasite clearance during the chronic rebound phase and PD-1 blockade, which is highly expressed on CD8^+^ T cells, further improved parasite elimination. With regards to CD4^+^ T cells, however, LAG-3 blockade most likely acts through unmasking CD4 interactions with MHC-II to ultimately promote higher secretion of effector cytokines and -possibly-more robust recruitment/activation of innate effector cells. Overall, these inhibitory pathways appear to slow down host clearance of the parasite in this non-lethal animal model of malaria; but further studies of these pathways in a lethal model to fully represent infection outcomes in humans, will be needed to understand the clinical implications of targeting these pathways during natural infection.

## Materials and Methods

### Mice

*This study was carried out in strict accordance with the recommendations by the animal use committee at the Albert Einstein College of Medicine that approved the animal protocol.*

Wild-type (WT) C57BL/6J (B6) 6-12 week-old male mice, PD-1^*-/-*^ and PD-L1^-/-^ (Gift Stan Nathenson, Einstein) were housed and bred in our SPF animal facility for all experiments.

### *Plasmodium* infections, blood parasitemia and adoptive serum transfers

#### Infections

*Plasmodium yoelii (Py) 17XNL(1.1)* parasites (stock MRA-593) and GFP expressing parasites (stock MRA-817) were obtained from the Malaria Research and Reference Reagent Resource Center as part of the BEI Resources Repository (NIAID, NIH, Manassas, VA. The *P. yoelii*, strain 17XNL(1.1), MRA-593 was contributed by D. J. Carucci and the *P. yoelii* strain 17XNL:PyGFP, MRA-817 was contributed by Ana Rodriguez. *Py*-infected red blood cells (iRBC) from a frozen stock (−80°C in Alsever’s solution, 10% glycerol) were intraperitoneally (i.p.) injected into a WT B6 mouse and grown for ∼4 days. When parasitemia reached 2-5%, 2×10^5^ *Py* iRBCs were injected intravenously (i.v.) into each experimental mouse.

#### Parasitemia and weight

Blood parasitemia was determined by flow cytometry on 1 µL of blood obtained by cutting the tip of the mouse tail with a sterile razor. Blood was fixed in 200µl of 0.025% glutaraldehyde in PBS 1mM EDTA before washing and permeabilization with 0.25% Triton X-100 in PBS for 5 min. After centrifugation, RBCs were incubated in 1mg/mL RNAse A (Sigma) for 30 min at room temperature (RT) and stained with 0.5 µM of the YOYO-1 dye (Invitrogen) for 30 min at RT and directly analyzed on a BD FACSCanto II (Becton Dickinson, CA). RBCs were gated based on forward and side scatter, and parasitemia was determined as the frequency of YOYO-1^+^ cells among all gated RBCs. Manual counting of Giemsa-stained blood smears by microscopy gave comparable parasitemia results. Mice were individually weighed (recorded in grams) on a tared weighing scale prior to blood collection.

#### Serum transfer experiments

Serum harvested from blood collected by cardiac puncture from indicated mice, at indicated time points was stored at -80°C. Serum was thawed, pooled within treatment groups and 150 µL of pooled serum were transferred to experimental mice intravenously (i.v.) prior to infection with *P. yoelii 17XNL(1.1)*. Pooled serum was heat inactivated at 56°C for 30 minutes prior to use in ELISAs.

### Antibody Blockade and depletion

#### PD-L1, PD-1 and LAG-3 blockade

200μg, unless otherwise indicated in figure legends, of polyclonal rat IgG (BioXcell; isotype) or anti-LAG-3 (clone C9B7W, BioXcell) and/or anti-PD-L1 (10F.9G2, BioXcell) or anti-PD-1 (RMP1-14, BioXcell) blocking Abs were injected i.v. into blood-stage parasitized mice beginning on day 9 or 12 after infection (as indicated) and every 3 days until day 15 or 18 as indicated. Antibodies which contained endotoxin levels above 3 EU/ml (Kinetic-QCL Kinetic Chromogenic LAL Assay, Lonza) were depleted of endotoxin with the ToxinEraser Endotoxin Removal kit (GenScript) following manufacturer’s instructions.

#### T cell depletions

150μg of anti-CD4 (clone GK1.5) or anti-CD8β (H35) mAbs were injected intraperitoneally (i.p.) one day before *P. yoelii 17XNL(1.1)* infection, and then at day 4 and 9 post infection.

### Preparation of cell suspensions for flow-cytometry (FACS) analysis

Spleens were dissociated on nylon meshes (100μm) and incubated at 37°C for 20 min in HBSS medium containing 4,000 U/ml of collagenase I (Gibco) and 0.1 mg/ml of DNase I (Roche), and RBCs further lysed with 0.83% NH_4_Cl buffer. Cells were resuspended in FACS buffer (PBS 1% FCS, 2mM EDTA, 0.02% sodium azide) and used for the different analyses detailed below.

### Cell staining for FACS analysis

Cell suspensions were incubated with 2.4G2 antibody for 15 min at 4°C and further stained with various antibody cocktails (S1 Table) in FACS buffer. For detection of intracellular IFNγ, TFNα and IL-2 staining, cells were incubated for 3-4 hrs at 37°C, 5% CO_2_ in RPMI 1640 (Invitrogen) 5% FCS, Golgi Plug (BD, 1/1000), Golgi Stop (BD, 1/1000), PMA (Fisher, 50ng/mL), Ionomycin (Fisher, 1μg/mL) and then fixed in IC fixation buffer (eBioscience) for 15 min at 4°C, and permeabilized for 30 min in 1X Perm/Wash buffer (eBioscience) containing indicated intracellular marker(s). For detection of transcription factors (BCL6, EOMES, FoxP3, T_bet_, TCF1), Ki67 and Granzyme B, cells were fixed and permeabilized after the surface stain with the FoxP3/Transcription Factor Staining Buffer Set following the manufacturer’s instructions (eBioscience). Stained cells were collected on BD LSR-II or Aria III and the Cytek Aurora. Data were analyzed using FlowJo version 9.6.6 or FlowJo version 10.5.3 (TriStar).

### ELISA for parasite antigens

Detection of antibodies against *P. yoelii* MSP1_19_ (BEI resources/ATCC, MRA-48) or *Py* iRBC antigens present in mouse sera collected at indicated time points, were done by ELISA following an adapted protocol (8). Briefly, sera dilutions were incubated for 1.5 hours at 37°C in high bind polystyrene microplate wells (Corning) that were first blocked with 1% BSA and then coated with either MSP1_19_ (2.5µg/ml) or *Py* iRBC lysate (50µg/ml). Total specific IgG sera antibodies were detected with horse radish peroxidase (HRP) conjugated goat anti-mouse IgG antibodies (Jackson ImmunoResearch) followed by addition of the substrate 3,3’,5,5’-tetramethylbenzidine (Sigma) and reading of absorbance at 450 nm. Endpoint titers were reported with the background absorbance of PBS coated wells subtracted. To obtain *Py* iRBC protein extracts, blood was collected by cardiac puncture from WT mice infected with *Py* (∼20-30% parasitemia) and placed in heparinized tubes. Blood was washed 3 times in RPMI media and centrifuged for 15 min at 1,500xg at 4°C. Pellets containing *Py*-iRBC were stored at -80°C before protein extraction. *Py* protein extract was obtained by adapting a previously described method (20). Briefly, frozen iRBC pellets were thawed and lysed directly in ultra-pure water containing protease inhibitor mix M (Roche) and 1mM PMSF (Sigma-Aldrich). This was followed by 4 washes with ice-cold PBS, 1mM PMSF, at 20,000xg for 25 min at 4°C and 3 freeze/thaw cycles in liquid nitrogen and 30 min room temperature incubations. Protein extracts were also obtained as described from RBC harvested from uninfected WT mice to use as negative controls for ELISA. Extracts from *Py* iRBCs and uninfected RBCs were stored at -80°C before utilization. Protein concentration of lysates were determined by Bradford Assay (Thermo Scientific).

### Statistics

Statistical significance was calculated using an unpaired Student t test with GraphPad Prism software and two-tailed P values are reported as: (*) P<0.05, (**) P<0.01, (***) P<0.001, (****) P<0.0001, (ns) P>0.1. All p values of 0.05 or less were considered significant and are referred to as such in the text.

## Acknowledgements

We thank the Einstein FACS and microscopy core facilities.

## Supporting Information

**S1 Table.**
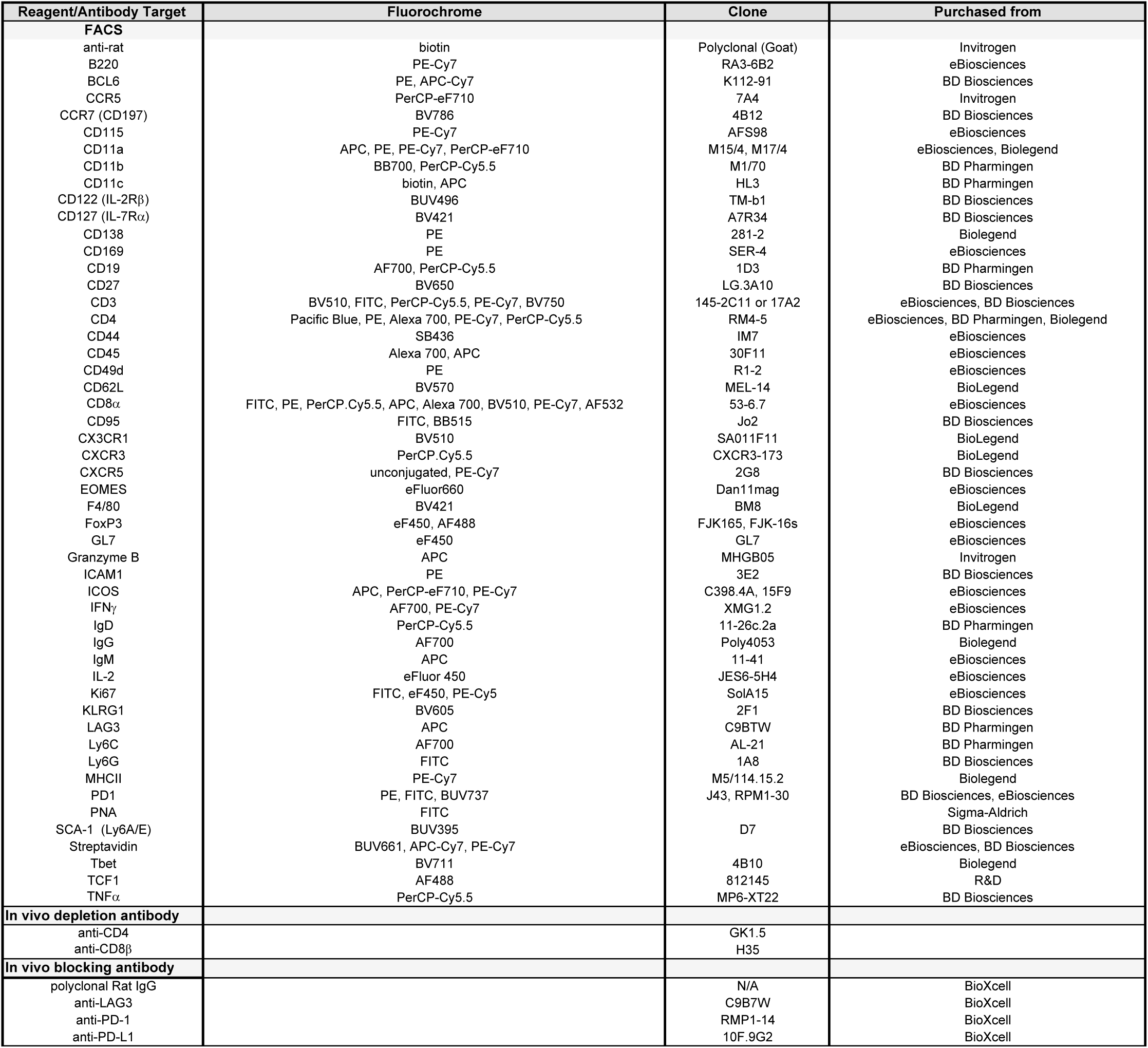
List of antibodies used for flow cytometry analyses and in vivo blocking or cell depletion.

| Reagent/Antibody Target | Fluorochrome | Clone | Purchased from |
| --- | --- | --- | --- |
| **FACS** |  |  |  |
| anti-rat | biotin | Polyclonal (Goat) | Invitrogen |
| B220 | PE-Cy7 | RA3-6B2 | eBiosciences |
| BCl6 | PE, APC-Cy7 | K112-91 | BD Biosciences |
| CCR5 | PerCP-eF710 | 7A4 | Invitrogen |
| CCR7 (CD197) | BV786 | 4B12 | BD Biosciences |
| CD115 | PE-Cy7 | AFS98 | eBiosciences |
| CD11a | APC, PE, PE-Cy7, PerCP-eF710 | M15/4, M17/4 | eBiosciences, Biolegend |
| CD11b | BB700, PerCP-Cy5.5 | M1/70 | BD Pharmingen |
| CD11c | biotin, APC | HL3 | BD Pharmingen |
| CD122 (IL-2Rβ) | BUV496 | TM-b1 | BD Biosciences |
| CD127 (IL-7Rα) | BV421 | A7R34 | BD Biosciences |
| CD138 | PE | 281-2 | Biolegend |
| CD169 | PE | SER-4 | eBiosciences |
| CD19 | AF700, PerCP-Cy5.5 | 1D3 | BD Pharmingen |
| CD27 | BV650 | LG.3A10 | BD Biosciences |
| CD3 | BV510, FITC, PerCP-Cy5.5, PE-Cy7, BV750 | 145-2C11 or 17A2 | eBiosciences, BD Biosciences |
| CD4 | Pacific Blue, PE, Alexa 700, PE-Cy7, PerCP-Cy5.5 | RM4-5 | eBiosciences, BD Pharmingen, Biolegend |
| CD44 | SB436 | IM7 | eBiosciences |
| CD45 | Alexa 700, APC | 30F11 | eBiosciences |
| CD49d | PE | R1-2 | eBiosciences |
| CD62L | BV570 | MEL-14 | BioLegend |
| CD8α | FITC, PE, PerCP-Cy5.5, APC, Alexa 700, BV510, PE-Cy7, AF532 | 53-6.7 | eBiosciences |
| CD95 | FITC, BB515 | Jo2 | BD Biosciences |
| CX3CR1 | BV510 | SA011F11 | BioLegend |
| CXCR3 | PerCP-Cy5.5 | CXCR3-173 | BioLegend |
| CXCR5 | unconjugated, PE-Cy7 | 2G8 | BD Biosciences |
| EOMES | eFluor660 | Dan11mag | eBiosciences |
| F4/80 | BV421 | BM8 | BioLegend |
| FoxP3 | eF450, AF488 | FJK165, FJK-16s | eBiosciences |
| GL7 | eF450 | GL7 | eBiosciences |
| Granzyme B | APC | MHGB05 | Invitrogen |
| ICAM1 | PE | 3E2 | BD Biosciences |
| ICOS | APC, PerCP-eF710, PE-Cy7 | C398.4A, 15F9 | eBiosciences |
| IFNγ | AF700, PE-Cy7 | XMG1.2 | eBiosciences |
| IgD | PerCP-Cy5.5 | 11-26c.2a | BD Pharmingen |
| IgG | AF700 | Poly4053 | Biolegend |
| IgM | APC | 11-41 | eBiosciences |
| IL-2 | eFluor 450 | JES6-5H4 | eBiosciences |
| Ki67 | FITC, eF450, PE-Cy5 | SolA15 | eBiosciences |
| KLRG1 | BV605 | 2F1 | BD Biosciences |
| LAG3 | APC | C9BTW | BD Pharmingen |
| Ly6C | AF700 | AL-21 | BD Pharmingen |
| Ly6G | FITC | 1A8 | BD Biosciences |
| MHCII | PE-Cy7 | M5/114.15.2 | Biolegend |
| PD1 | PE, FITC, BUV737 | J43, RPM1-30 | BD Biosciences, eBiosciences |
| PNA | FITC |  | Sigma-Aldrich |
| SCA-1 (Ly6A/E) | BUV395 | D7 | BD Biosciences |
| Streptavidin | BUV661, APC-Cy7, PE-Cy7 |  | eBiosciences, BD Biosciences |
| Tbet | BV711 | 4B10 | Biolegend |
| TCF1 | AF488 | 812145 | R&D |
| TNFα | PerCP-Cy5.5 | MP6-XT22 | BD Biosciences |
| **In vivo depletion antibody** |  |  |  |
| anti-CD4 |  | GK1.5 |  |
| anti-CD8β | | H35 | |
| **In vivo blocking antibody** |  |  |  |
| polyclonal Rat IgG |  | N/A | BioXcell |
| anti-LAG3 |  | C9B7W | BioXcell |
| anti-PD-1 |  | RMP1-14 | BioXcell |
| anti-PD-L1 |  | 10F.9G2 | BioXcell |

**S1 Fig.**
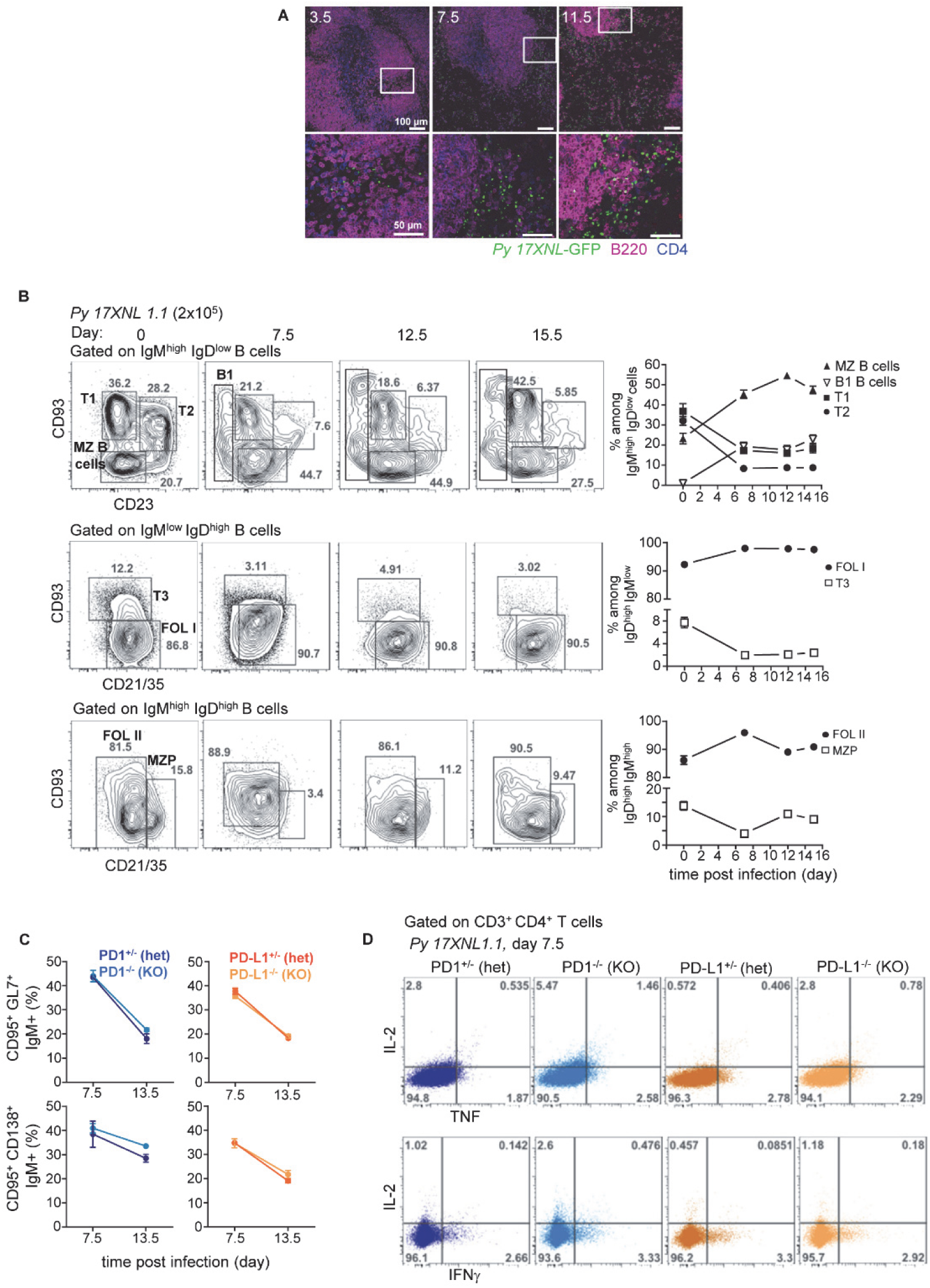
B cell and CD4^+^ T cell responses in the spleen of *Py*-infected mice. **(A)** At indicated days after 2×10^5^ *Py*GFP^+^ inoculation, spleens were fixed overnight in 1% PAF in PBS, washed in phosphate buffer, and dehydrated in 30% sucrose in phosphate buffer for 12 hrs. Spleens were snap frozen in TissueTekH (Sakura Finetek) and cut into 10 mm sections prior staining with an anti-B220 mAb (clone RA3-6B2, BD) and amplified with an anti-rat Alexa Fluor 568. Sections were further stained with anti-CD4 (clone RM4-5, BD) and anti-GFP to detect GFP expressing parasites and amplified with an anti-rat Alexa fluor 633 and anti-chicken Alexa Fluor 488 respectively. Images were acquired on a SP5 confocal microscope (Leica) and processed using ImageJ Adobe Photoshop software. **(B)** Spleens from *Py*-infected mice were harvested at indicated days post infection and cells stained with mAb against lineage markers (CD19, B220) and activation/subset markers (IgD, IgM, CD93, CD23, CD21/35, GL7, CD138, CD95, PNA). In all experiments, representative FACS dot plots of 3-5 independent replicate experiments are presented for 1 of 5-18 mice. Graphs average each experiment shown with SEM. P-values are indicated when applicable. (C, D) Mice of indicated genotypes were inoculated with 2×10^5^ *Py* 17XNL1.1 iRBC i.v. **(C)** Spleens were harvested at indicated days post infection and cells stained with mAb against lineage markers (CD19, B220) and activation/subset markers (IgD, IgM, CD93, CD23, CD21/35, GL7, CD138, CD95, PNA). Graphs average 3-5 independent experiment with SEM (n=5=18). **(D)** As in Fig 3, spleen cells from 7.5 *Py*-infected littermates mice of indicated genotypes were incubated with PMA/ionomycin for 4 hours and stained for cell-surface CD3 and CD4 and intracellular cytokines IL-2, IFNγ and TNF. Frequencies of cytokine-producing cells among CD4^+^ T cells are shown in a representative FACS dot plots of 2 independent replicate experiments are presented (n= 3-12).

**S2 Fig.**
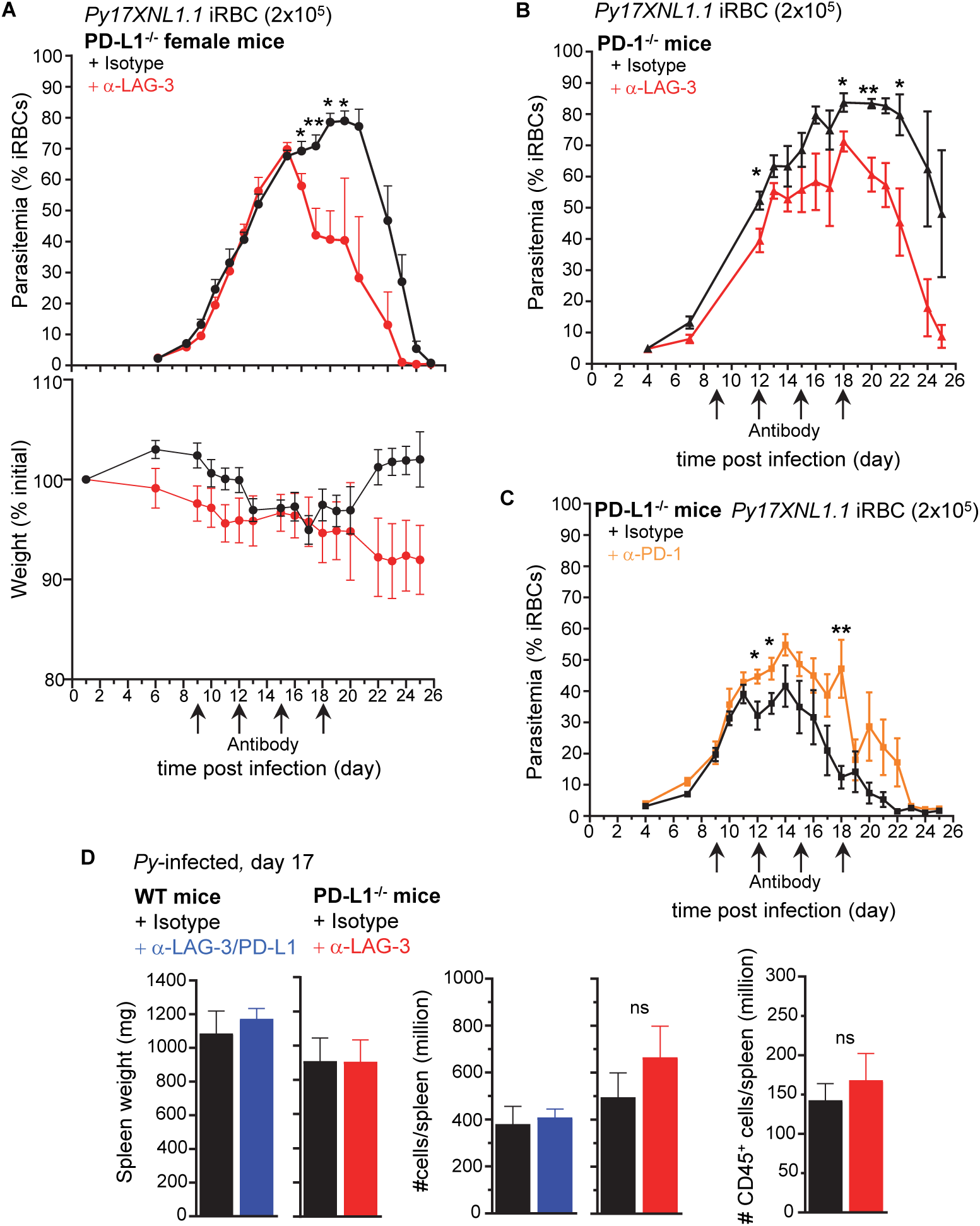
LAG-3 blockade in female PD-L1^-/-^ mice and PD-1^-/-^ mice promotes reduced parasitemia. **(A-C)** Indicated genotype and gender mice were inoculated with 2×10^5^ *Py* 17XNL1.1 iRBC i.v., and 9, 12, 15 and 18 days post infection, indicated mice received 300 μg (A) or 200 μg (B-C) of anti-LAG-3 or anti-PD-1 or isotype Ab i.v. Results show the kinetics of blood parasitemia over time, measured starting day 4 post infection and every day or every other day until day 25 using YOYO-1 staining of RBC and FACS. Graphs average the pool of 1-2 independent experiments shown with SEM (n=4-10). P-values are indicated when applicable (*p<0.05, **p<0.01) for each treatment compared to isotype group per time point. **(D)** Spleen weight, total and hematopoietic (CD45^+^) cell numbers in WT or PD-L1^-/-^ mice treated (day 9, 12, 15) with anti-LAG-3/PD-L1, anti-LAG-3 or matching isotype 17 days post *Py* infection pooled across 2 experiments are shown (n=4-10 mice).

**S3 Fig.**
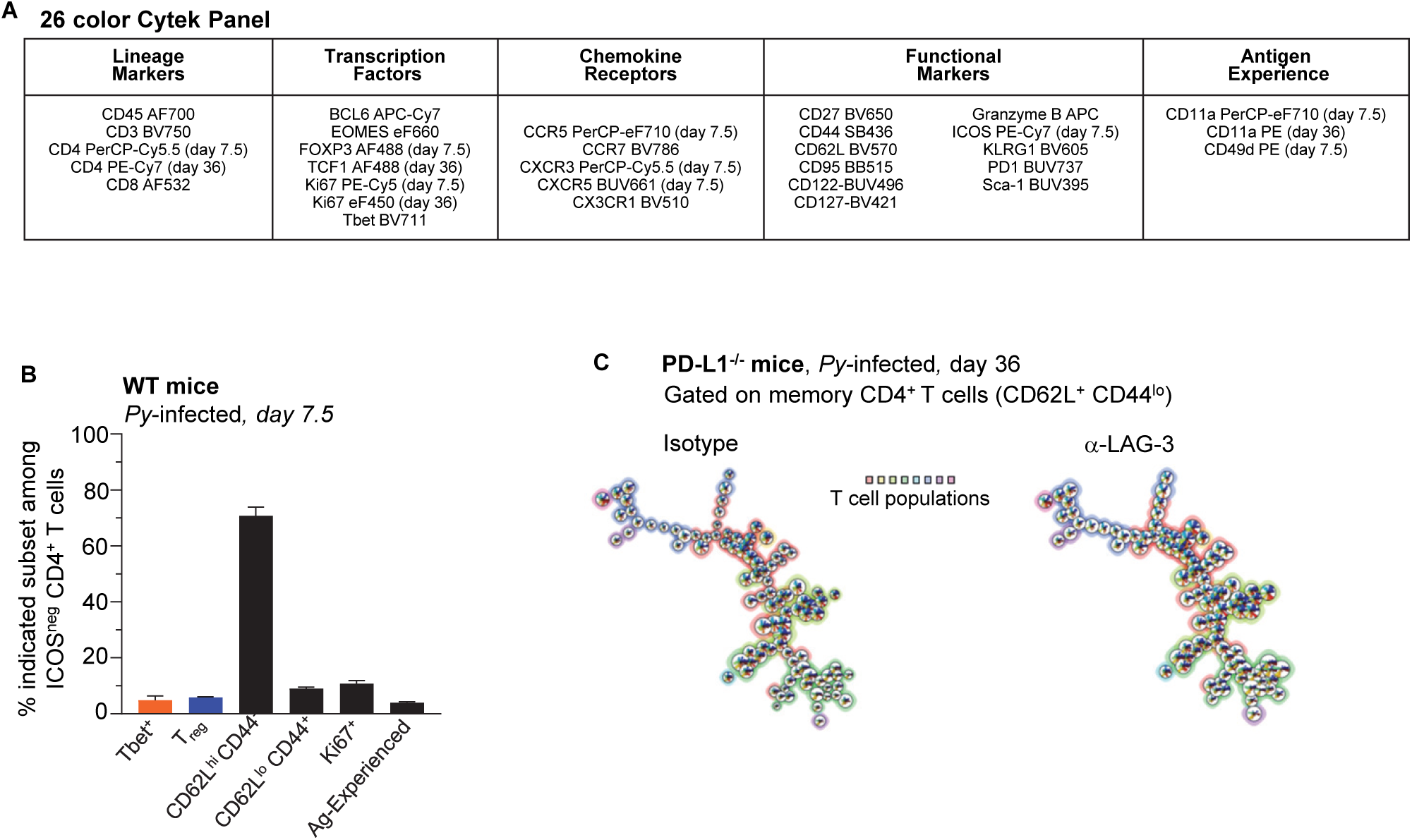
High dimensional flow cytometry panel to characterize CD4^+^ and CD8^+^ T cells in *Py-*infected mice. (A) Panel of Abs applied to splenocytes from WT mice at either day 7.5 or day 36 post *Py 17XNL1.1* infection, as indicated, for staining and analysis on the Cytek Aurora cytometer. **(B)** Indicated cell subset frequencies among ICOS negative splenocytes in WT mice on day 7.5 post *Py* infection. **(C)** As in Fig 9D, Spleen cells from day 36 PD-L1^-/-^ *Py*-infected mice treated with either anti-LAG-3 or isotype Ab i.v. (on days 9, 12, 15) were stained for lineage markers (CD8, CD4, CD3, Bcl6, T_bet_) and functional markers (CD62L, CD44, CD11a) with a 24 color panel depicted in (A). FlowSOM analysis on pooled (n=4-5) memory (CD62L^+^ CD44^lo^) CD4^+^ T cells are presented.

**S4 Fig.**
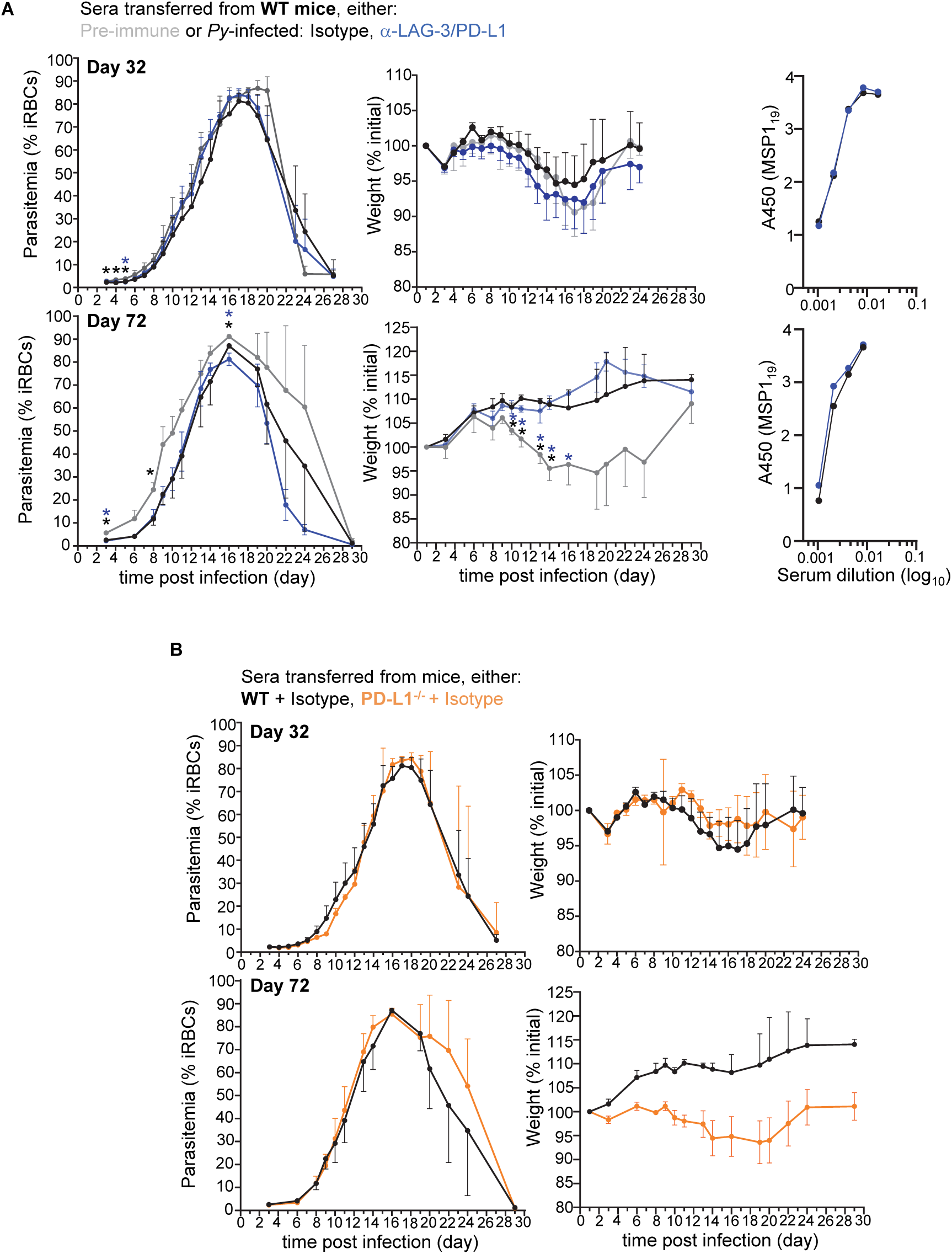
LAG-3 neutralization in WT mice fails to promote the production of higher titers and more protective parasite-specific antibodies. Blood parasitemia and change in initial weight of WT mice measured over time starting day 3 post inoculation with 2×10^5^ *Py* 17XNL1.1 iRBC i.v. Prior to infection, mice received 150 µl of sera from **(A)** either pre-immune WT mice or sera collected from isotype treated or α-LAG-3/PD-L1 treated *Py-*infected WT on either day 32 or 72 post infection. Titers of MSP_1-19_ –specific IgG Abs in indicated sera detected by ELISA as described in the methods. In **(B)** sera comes from either isotype treated *Py-*infected WT mice or *Py-*infected PD-L1^-/-^ mice at day 32 or 72.

